# Drought alters volatile profiles in European beech saplings across genetically diverse backgrounds

**DOI:** 10.64898/2026.06.24.733805

**Authors:** Ting Tang, Toja Guerra, Domitille Coq-Etchegaray, Bernhard Schmid, Linus Reichert, Guido L. B. Wiesenberg, Meredith C. Schuman, Sofia van Moorsel

## Abstract

- European beech (*Fagus sylvatica* L.) is a widely distributed, ecologically and economically important deciduous tree species in European forests, but is increasingly threatened by drought stress. Volatile organic compounds (VOCs) are ubiquitous plant metabolites that may serve as non-invasive biomarkers of drought stress, yet they have rarely been studied in European beech.
- In this study, we examined VOC responses of European beech to experimental drought across diverse genetic backgrounds in a common garden. The 72 four-year-old beech saplings represented three genetic clusters, seven provenances (geographic seed sources), and 12 maternal seed families. Half of the saplings were assigned to the drought treatment and received no water for 14 days, while the remaining saplings served as controls and were watered as required. VOC profiles, quantified as peak heights of mass spectral features, were measured for all individuals during pre-drought, drought, and rewatering periods.
- We found that pre-drought VOC profiles, in particular monoterpenes, varied significantly among genetic backgrounds. Experimental drought significantly altered VOC profiles, characterized by increased green leaf volatiles and decreased monoterpenes, oxidized terpenoid derivatives, and other fatty acid derivatives. Reductions in monoterpenes persisted after rewatering, indicating a drought legacy effect. Drought responses were largely conserved across genetic backgrounds, with significant seed family-specific responses detected for only three VOC features.
- Our findings suggest that VOC profiles are genetically structured yet highly plastic under drought and highlight their potential as non-invasive biomarkers for monitoring drought stress in European beech under climate change.

## Introduction

European beech (*Fagus sylvatica* L.) is a widely distributed deciduous tree species characteristic of many forests across Europe (Durrant et al., 2016; Fuchs et al., 2024). It is ecologically important because of providing unique habitat for diverse wildlife and of economic importance due to its varied applications in forestry(Durrant et al., 2016). However, climate change exposes European beech to more frequent and intensive drought and threatens its continued occurrence across the continent (Martinez del Castillo et al., 2022; Antonucci et al., 2021).

European beech exhibits substantial genetic variation that reflects phylogeography and aligns with its geographical distribution (Lazic et al., 2024; Milesi et al., 2024; Stefanini et al., 2022). Studies have suggested that morphological and physiological traits of European beech under drought vary by geographic origin (*i.e*., provenance) (Dounavi et al., 2016; Knutzen et al., 2015; Pšidová et al., 2015). These findings indicate that genetic differences in drought responses may have developed under local environmental conditions. Other studies report substantial within-provenance variation in drought responses, driven by within-provenance genetic and phenotypic variation (Aranda et al., 2017; Bresson et al., 2011; Wortemann et al., 2011). This may be reflected in differences among maternal seed families (half-to full-sibs depending on out-crossing rate) within provenance. In addition to genetic variation, phenotypic plasticity allows a single genotype to produce different phenotypes in response to environmental variation (Sultan, 1995). Genetic variation and phenotypic plasticity have been shown to shape drought-related traits in European beech—such as growth-related (Frank et al., 2017), morphological (Stojnić et al., 2015), root (Meier and Leuschner, 2008), and physiological traits (Sánchez-Gómez and Aranda, 2024). Despite extensive research on drought impacts on European beech across diverse genetic background (reviewed by Leuschner, 2020), we still lack reliable early biomarkers for detecting drought stress in individual trees.

Plant volatile organic compounds (VOCs) are metabolites ubiquitous in within-plant and between-plant signaling, as well as interactions with pollinators, microorganisms, and seed dispersers (Baldwin, 2010). Plants also produce VOCs in response to biotic stress, such as herbivore attack and pathogen infection, and abiotic stress such as drought and heat (reviewed by Niederbacher et al., 2015; Loreto and Schnitzler, 2010; Schuman, 2023). Given their stress-responsive nature, VOCs can serve as efficient, non-invasive airborne biomarkers of plant responses to drought as they evaporate at normal temperatures and can be sampled from headspace air (Tholl et al., 2021).

Drought stress can affect VOC emissions by reducing stomatal conductance and photosynthesis, causing oxidative stress and membrane damage (Farooq et al., 2009; Hasanuzzaman et al., 2013). Different classes of VOCs play different roles and respond differently to stress depending on their biosynthetic pathways and specific regulation (reviewed by Schuman, 2023). Terpenes and green leaf volatiles (GLVs) are among the most common VOCs involved in stress responses. Terpene biosynthesis depends on photosynthetic carbon, but terpene emissions may continue under mild drought by using stored carbon sources such as starch, whereas prolonged or severe drought can reduce VOC production due to limited carbon and energy availability (Loreto and Schnitzler, 2010; Possell and Loreto, 2013). Some terpenes, such as isoprenes and monoterpenes, can stabilize cell membranes and reduce oxidative stress (Vickers et al., 2009; Loreto and Schnitzler, 2010). GLVs are rapidly emitted after tissue damage and are well studied in biotic stress responses. They can directly attract or repel herbivores, act as damage signals that prime or induce defenses in undamaged tissues and neighboring plants, and attract natural enemies of herbivores, thereby enhancing indirect plant defense (Engelberth et al., 2004; Ameye et al., 2018; Matsui and Engelberth, 2022). More recent studies suggest that GLVs may also respond to drought stress (Catola et al., 2016; Jin et al., 2023; Shen et al., 2026). After drought stress, plant VOC emissions generally recover toward pre-drought levels as photosynthesis resumes following rewatering (Šimpraga et al., 2011; Staudt et al., 2008). However, VOC recovery is often delayed relative to photosynthesis and may remain incomplete after rewatering, especially after severe drought (Männistö et al., 2024). Yet, VOC responses to drought are mostly documented in herbaceous (Reinecke et al., 2024; Campbell et al., 2019) and crop species (Weldegergis et al., 2015; Pagadala Damodaram et al., 2021), with a few largely cultivated trees like tea and fruit trees (Catola et al., 2016; Jin et al., 2023). Forest trees, and European beech in particular, remain understudied despite the urgent need to understand their drought responses.

In this study, we investigated how drought stress affects VOC profiles in European beech across multiple hierarchical levels of genetic variation (referred to as “genetic background”). We used 72 four-year-old European beech saplings representing 12 maternal seed families (seeds collected from a single tree or very close (< 10 m) to it) nested in seven provenances and these nested in three genetic clusters (Fig. 1a). Half of the saplings (as far as possible within each seed family) were subjected to a drought treatment and the other half to a control treatment. VOC samples were collected during three periods: before drought (pre-drought), during drought, and after rewatering (Fig. 1c). First, we examined how genetic background influenced European beech VOC profiles under non-stress conditions (pre-drought period). We hypothesized that genetic background explains significant proportions of VOC variation among genetic clusters, provenances within clusters, and seed families within provenances (Fig. 1a). Furthermore, we expected that the proportion of VOC variation explained by genetic factors would differ among VOC classes. Second, we tested how drought affected the VOC profile of European beech by using the pre-drought period as a baseline (Fig. 1b). Specifically, we compared VOC peak heights during the drought and rewatering periods relative to pre-drought levels and examined the effects of treatment (drought vs. control) and its interaction with genetic background on VOC changes. We hypothesized that saplings subjected to drought would exhibit changes in VOC emissions relative to the control treatment from the pre-drought to the drought period (direct drought effect) and from the pre-drought to the re-water period (drought legacy effect, Fig. 1b). Given the common occurrence of genotype-by-environment interactions, we also expected that VOC responses to drought would depend on seed family background, which includes genetic variation among genetic clusters and provenances. However, due to the limited knowledge of VOC responses to drought in European beech and the complexity of VOC profiles, we did not make specific predictions about the direction of these differences.

**Figure 1.**
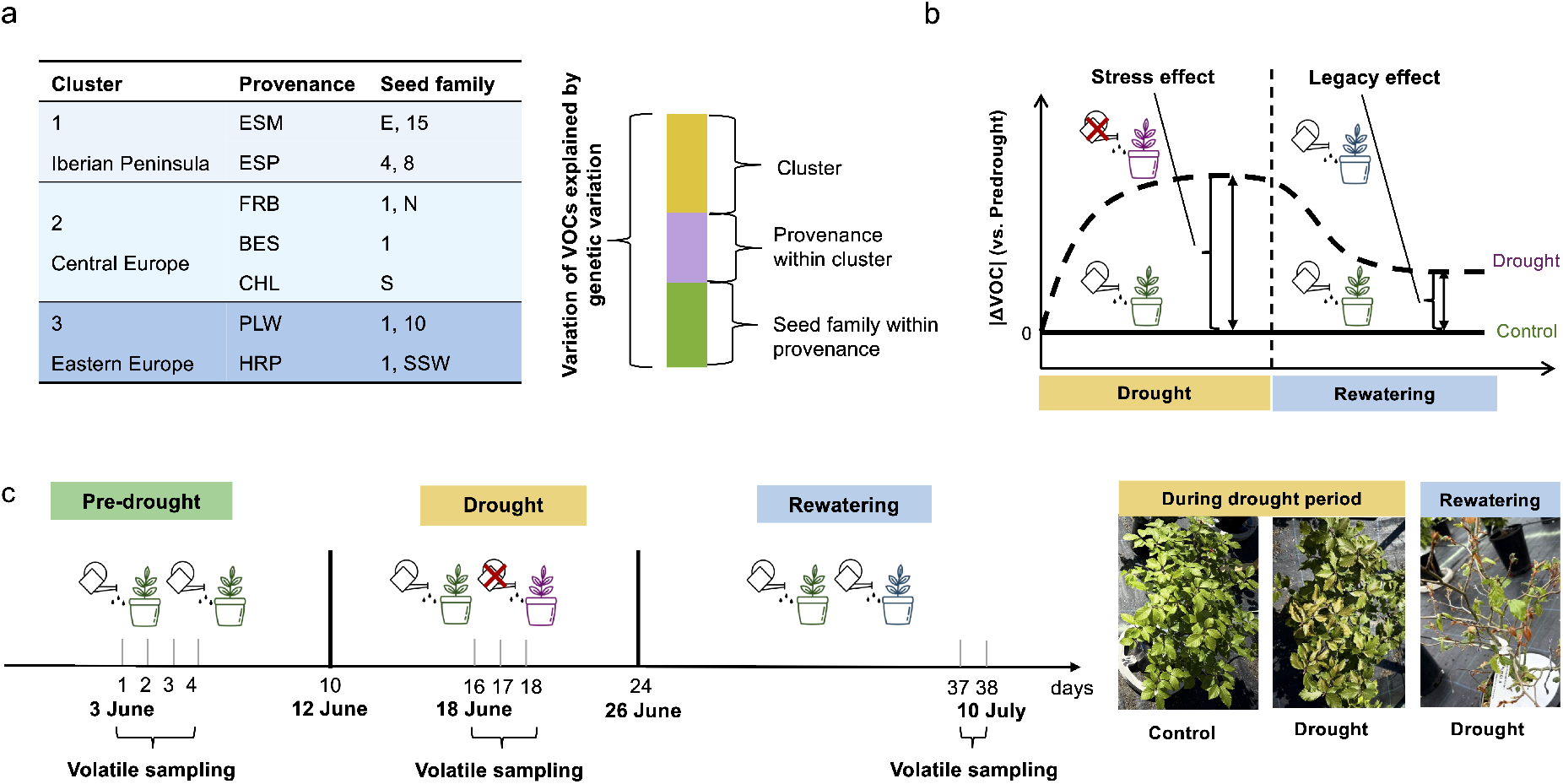
Conceptual framework and sampling timeline of the study. **(a)** Conceptual illustration of VOC variation explained by genetic background at three hierarchical levels: genetic cluster, provenance, and seed family. We hypothesized that VOC variation of European beech under non-drought conditions differed among the entities at these three genetic levels. Genetic clusters were inferred from whole-genome sequencing data; provenances indicate the geographical origins. Trees grouped within a seed family are expected to share the same maternal origin(s) based on seed collection records (numbers: same mother tree; letters: same location on ground, E: east, N: north, S: south, SSW: south to southwest, see text). **(b)** Illustration of drought-drought and drought-legacy effects. VOC peak heights during the drought and rewatering periods were compared to pre-drought levels, and the effect of treatment (drought vs. control) was tested. The differences between drought and control treatments during the drought period indicate a stress effect, whereas differences during the rewatering period indicate a drought-legacy effect. **(c)** Experimental timeline showing VOC sampling dates and the duration of the drought treatment. Photographs illustrate visible drought stress and recovery symptoms during the drought period and during rewatering, respectively.

## Materials and Methods

### Experimental design

This study was conducted in a common garden at the University of Zurich, Zurich, Switzerland (47^°^ 23^′^ 44^′′^ N, 8^°^ 33^′^ 05^′′^ E; 509 m a.s.l.). The common garden consists of 180 four-year-old European beech saplings from 16 provenances across Europe. For this study, we used a subset of 72 saplings from seven provenances (geographic seed source). Saplings were derived from seeds collected directly from the same mother tree or, when insufficient seeds were available (four of the 12 maternal groups used in this study), from one location likely representing the footprint of a single maternal tree (< 10 m radius) on the forest ground (see Czyż et al., 2025). Although these four maternal groups might sometimes have included seeds from maternal different trees, we here still call them maternal seed families (Fig. 1a). At the time of the experiment, the saplings were growing outside in 7-liter pots filled with an equal mixture of peat-free potting soil and low-organic sandy soil. For details on seed collection, germination, and garden establishment, see Kurath et al., 2025.

Based on whole-genome sequencing data, three genetic clusters were identified among the saplings in the common garden (Kurath et al., 2025). These clusters corresponded to the Iberian Peninsula (Cluster 1), Central Europe (Cluster 2), and Eastern Europe (Cluster 3), consistent with patterns reported in other pan-European genomic studies of European beech (Lazic et al., 2024; Milesi et al., 2024). From each genetic cluster, we selected two or three provenances and from each provenance one or two seed families depending on availability, resulting in four seed families per cluster (Supporting Information Fig. S1a). Each seed family was represented by six individual saplings (3 clusters × 4 seed families × 6 saplings; Supporting Information Fig. S1a).

Within each seed family, three saplings were randomly assigned to the drought and three to the control treatment, resulting in 36 saplings per treatment. The treatments were applied at block level, with nine saplings in each of eight blocks, four receiving the drought and four the control treatment (Supporting Information Fig. S1b). Within each drought block, each genetic cluster was represented by three saplings and all nine saplings belonged to different seed families, but otherwise the saplings were randomly selected with regard to provenance and seed-family identity. The positions of the selected saplings within each drought block were also randomized. Each control block was placed next to a corresponding drought block.

During the experiment, the saplings were irrigated by an automated micro-drip system (Micro-Drip-System, Gardena, Ulm, Germany). Saplings from the control treatment, as well as all saplings during the pre-drought and rewatering periods (Fig. 1c), were watered twice a day using the irrigation system at 6:30 a.m. and 5:00 p.m., for 10 minutes each. At the start of the drought period, cone-shaped rain covers were placed around the stems above the pots to exclude rainfall for both drought and control treatments (Kurath et al., 2025). The rain covers were made from waterproof and UV-resistant PVC black pond foil (Heissner GmbH, Lauterbach, Hessen, Germany) and fixed on the sapling stem with Parafilm (Pechiney Plastic Packaging Inc., Chicago, IL, United States) beneath the first stem node. During the drought period, the irrigation system was removed from the pots in the drought treatment. During rewatering, we removed the rain cover for all the saplings and reinstalled the irrigation system for the drought treatment. The drought treatment lasted 14 days. The detailed experimental timeline can be found in Fig. 1c.

To monitor soil conditions, 20 soil data loggers, each equipped with one moisture sensor and three temperature sensors (TMS-4, TOMST, Prague, Czech Republic; Wild et al., 2019), were installed in the pots of randomly selected drought-treated and control saplings (10 each). The loggers recorded soil moisture below the soil surface down to 14 cm and temperature at three soil depths (+15, 0 and −8 cm from the soil surface) every 15 minutes. Additionally, an all-in-one weather station (ATMOS 41, METER Group, Pullman, WA, USA) was installed in the same experimental field to record precipitation and air temperature every 10 minutes.

### VOC sampling

Volatile organic compounds (VOC) were sampled three times from all 72 saplings: once before the drought treatment, once during the drought treatment, and once during rewatering (Fig. 1c). Samples were collected during daylight hours between 9:30 and 17:00. Due to the required VOC collection time and time required for set-up and take-down, the 72 saplings could not be sampled simultaneously within a single day. To minimize temporal effects on plant VOC production (Chen et al., 2020) and to make sure that time of the day was not confounded with treatment, two paired blocks were always sampled in parallel, one from the drought and one from the control treatment (A and E; B and F; G and C; or D and H, see Supporting Information Fig. S1b) within a half day (approximately four hours). With the randomization of trees with different genetic backgrounds (see Experiment design) and the paired sampling of a drought and a control block, variation in VOC emission due to sampling time does not bias the effects we are interested in (i.e. genetic background and drought treatment). Each sampling period includes four half-day sessions, but the sampling schedule was adjusted to avoid days with precipitation, resulting in four sampling days for the pre-drought period, three sampling days for the during-drought period, and two sampling days for the rewatering period. During the drought period, VOCs were collected once some drought-treated saplings began to show visible drought stress symptoms, such as leaf yellowing (Fig. 1c). During rewatering, before sampling VOCs, we waited 13 days to allow the drought-treated saplings to show signs of recovery, such as newly flushed leaves (Fig. 1c). The full sampling timeline is shown in Fig. 1c.

A “push–pull” system was used to sample headspace VOCs from whole saplings enclosed in nylon bags, with glass tubes packed with Tenax^(R)^ TA (35/60 mesh, C1-BXXX-5039, Markes International, Llantrisant, UK; Supporting Information Fig. S1c) used to trap VOCs. The specific Tenax masses and recipes are proprietary and thus not provided by the manufacture. A 122 × 76.2 cm nylon oven bag (approximate diameter 48.5 cm; Xiamen Threestone Packing Material, Fujian, China) was supported by a wooden frame and secured with a flattened metal clothes hanger over the saplings (Supporting Information Fig. S1c). Nylon was selected as the enclosure material because it has low adsorption for plant-emitted VOCs (Morris et al., 2024) and its background signals are distinguishable from plant-derived VOCs. In addition, the flexible, large-volume bag ensured an approximately consistent headspace volume across saplings of varying sizes. Nylon bags were conditioned in an oven at 150 ^°^C for one hour per side (inside and outside) to remove VOCs from the bag material prior to use.

Because bagging can increase heat stress in plants, a fan-based “push” system was designed to reduce plant stress and enhance VOC collection efficiency. A hole (ca. 40 × 40 mm) was cut at the base of the sealed bag, and a 40 × 40 × 10 mm computer fan (NF-A4x10, Noctua, Vienna, Austria) was magnetically attached to the bag frame at the position of the cut hole, with the setup resting on the pot without direct contact. The fan was mounted in a customized 3D-printed polylactic acid (PLA) frame, which also held a sheet of activated charcoal filter positioned in front of—but not touching—the fan to filter incoming air. The fan operated at 5.0 V, producing an airflow of 0.17–0.20 m s^−1^, as measured in the laboratory using a thermal anemometer (testo 405i, Testo AG, Mönchaltorf, Switzerland).

The “pull” component of the system consists of an exhaust outlet and a Tenax^(R)^ TA tube VOC sampling setup. First, an outlet hole (ca. 2 × 2 cm) was cut at the top of the bag to create directed airflow across the plant, from the air inlet at the base to the outlet at the top (Supporting Information Fig. S1c). This hole was kept open by the metal clothes hanger, which also supported the bag on the frame (Supporting Information Fig. S1c). VOCs were sampled actively using programmable pump systems operating at a flow rate of 300 mL min^−1^ for 60 minutes. The pump system was adapted from the design of Geckeler et al., 2023, replacing the original control system with a programmable control system, integrating closed-loop flow control using an flow sensor (Omron D6F-01A1-110), increasing battery capacity, and adding WiFi-based remote control and firmware update capabilities. The airflow through the Tenax^(R)^ TA tubes was tested and verified using a flow meter (red-y compact, Vögtlin Instruments GmbH, Switzerland). The Tenax^(R)^ TA tube was inserted through the bag surface near the top and secured in place with a clip (Supporting Information Figure S1c). Before sampling, Tenax^(R)^ TA tubes were conditioned at 210 ^°^C for 30 mins in either a Gerstel TC2 Tube Conditioner (Heritage ALT LLC, USA) or at 220 ^°^C for 20 minutes with conditioning flow 80 mL min^−1^ using a thermal desorption autosampler attached to a gas chromatograph coupled to an electron impact ionization single quadrupole mass spectrometer (TD-GC-EI-MS). Matter desorbed during tube conditioning was not analyzed.

After sample collection, the two sides of each Tenax^(R)^ TA tube were wrapped immediately in polytetrafluoroethylene (PTFE) tape to prevent contamination during transport and storage and were kept tightly wrapped and away from strong odors until they were analyzed. Once the Tenax^(R)^ TA tubes were transported in the lab, they were stored at 4 ^°^C. in a refrigerator used only for non-volatile substances or trapped VOC samples. VOCs trapped on Tenax^(R)^ TA were extracted and analyzed by TD-GC-EI-MS.

TD-GC-EI-MS was performed using a Nexis GC-2030 coupled to a QP2020 NX EI-MS and a TD-30R thermal desorbtion unit (Shimadzu Corporation, Kyoto, Japan). Desorption was conducted at 220 ^°^C with nitrogen (purity 4.5) at 80 mL min^−1^. Volatiles were collected on a Tenax^(R)^ TA cold trap maintained at -20 ^°^C. Rapid cold trap desorption was carried out for 4 min at 230 ^°^C, with the valve, joint, and transfer line all held at 230 ^°^C. GC injection was done at a split ratio of 5:1 onto an RTX-WAX column (Restek, USA; i.d. × 0.25 µm film thickness) using helium as carrier gas (purity 5.0; PanGas, Dagmersellen, Switzerland) at a constant linear velocity of 40 cm s^−1^. The oven temperature program was: 40 ^°^C for 2 min; ramped 10 ^°^C min^−1^ to 150 ^°^C and held for 2 min; ramped 3 ^°^C min^−1^ to 190 ^°^C; ramped 10 ^°^C min^−1^ to 230 ^°^C; and held at 230 ^°^C for 3 min. Electron impact ionization was done at 70 eV, with the ion source and interface temperatures set to 230 ^°^C. The single-quadrupole mass analyzer scanned a mass range of m/z 33–400 from 3 to 30 min at 5 scans *s*^−1^. Data were recorded as m/z with ion counts as intensity.

We included one glass blank (a glass tube without Tenax TA sorbent) for each TD–GC-EI–MS analysis. In addition, we measured four types of blank controls for each treatment of 18 saplings sampled at the same time: (1) Setup controls: empty bags above three pots containing the same soil mixture and an identical sampling setup but no sapling (Supporting Information Fig. S1); (2) Ambient air controls: three ambient air samples collected at the field site near the potted saplings; (3) Field blanks: three clean Tenax^(R)^ TA tubes transported to the field but not used for sampling; and (4) Lab blanks: three clean Tenax^(R)^ TA tubes kept in the laboratory.

### Data analyses

#### Soil moisture and temperature

To verify the effectiveness of the drought treatment, soil moisture and temperature data were extracted from soil data loggers for the period from 28 May 2025 to 12 July 2025. We calculated daily mean values and standard errors of soil moisture, as well as mean soil temperature for the three sensing depths. Precipitation and air temperature data were extracted for the same period. Daily total precipitation and the daily mean and standard error of air temperature were calculated as additional references to complement the soil logger data.

#### VOC data processing

##### Raw VOC data processing

Raw TD–GC-EI–MS data were converted to the open format mzXML and processed in MZmine 4.7.27 (Schmid et al., 2023). The workflow included mass detection, extracted ion chromatogram (EIC) construction, chromatographic peak deconvolution, and retention time alignment across samples. Compound annotation was performed using the NIST 20 mass spectral library, with a minimum cosine similarity threshold of 85%. The parameter settings in MZmine followed the published MZmine protocol and were further optimized based on expert knowledge for this dataset. All parameters were implemented using batch mode, and the MZmine batch file is available on Dryad repository (see Data availability).

For each detected mass spectral feature (hereafter referred to as VOC feature), peak height, m/z ratio, retention time, and tentative annotation were exported for downstream analyses. All detected features were subsequently manually inspected in Shimadzu Postrun Analysis (version 4.6 SP1) software to confirm peak detection and annotation. In cases where no annotation was assigned in MZmine but a clear chromatographic peak was detected in Postrun and the library match exceeded 85%, the Postrun annotation was used for compound classification. Based on the compound tentative annotations, VOCs were classified according to their biosynthetic pathways following Supporting Information Table S4 which reformatted from Table S1 in Schuman, 2023.

##### Blank subtraction

The VOC features mean peak height detected in glass blanks were quantitatively removed from all tree VOC samples and the other blank controls. For each sample period, the mean peak height of the corresponding setup controls, ambient air, field and lab blanks were removed from the tree VOC samples. If subtraction of the tree VOC peak height resulted in a negative value, the feature was considered undetected and its peak height was set to zero.

##### Rare features filtering

To remove features uninformative for determining statistical patterns, we excluded features present in fewer than 18 samples (*i.e*., half of the trees in either the control or drought treatment). We conducted two types of analyses: (1) across the three sampling periods to evaluate changes in VOC profiles over the time of drought treatment and rewatering, and (2) within each sampling period to compare the drought and control treatments.

For the time-scale analysis, VOC features were filtered across all three sampling periods combined. Specifically, a feature was retained if it was detected in at least 18 out of the total 216 samples (72 trees × 3 sampling periods). For analyses conducted separately within each sampling period, filtering was applied independently for each period. In this case, only features detected in at least 18 out of 72 samples for each period were retained.

##### Feature detection and classification

After the filtering for blanks and rare features, we found 57 features across 72 trees sampled at three time points (pre-drought, drought, and rewatering). We classified the features into 11 volatile classes nested in six biosynthetic treatments (Schuman, 2023, Supporting Information Table S4 & S1). The tentative annotation and classification of each feature are provided in Supporting Information Table S3. Features identified as contaminants were excluded from the statistical analyses.

#### Statistical analyses

Data processing after mzmine and all other statistical analyses were performed in R 4.3.3 (www.r-project.org). All LMMs were fitted by the “lmer” function of the “lme4” package (version 1.1.37, Bates et al., 2015) and “lmerTest” was used to assess statistical significance unless otherwise specified. Peak height was *log*_10_-transformed (log_10_(*Peak_Height*_*ijk*_ + 1)) to achieve consistent error variance and normally distributed residuals; the addition of 1 was necessary to retain observations that were zero at the original scale.

##### Genetic effects

To test the overall effect of genetic background on VOC variation (H1), we used VOC data from all 72 trees collected before the drought treatment was applied to fit a linear mixed-effects model (LMM). In the overall model, *log*_10_-transformed peak height was specified as the dependent variable, while cluster, provenance, and seed family, as well as VOC class and its interaction with genetic factors, were included as fixed effects. Feature ID was included as a random factor. Block was not included as a random factor because genetic backgrounds (genetic cluster, provenance, and seed family) were randomized across blocks, making block and genetic background orthogonal (see Experiment design):

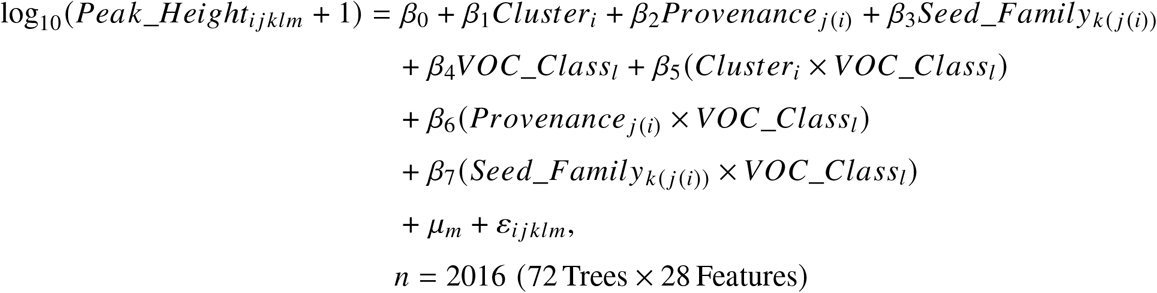

where log_10_(*Peak_Height*_*ijklm*_ is the *log*_10_-transformed peak height of VOC class *l* measured in seed family *k* nested within provenance *j* and cluster *i*, and feature *m*; *β*_0_ is the intercept; *β*_1_*Cluster*_*i*_, *β*_2_*Provenance*_*j*(*i*)_, and *β*_3_*Seed_Family*_*k*(*j*(*i*))_ are the fixed effects of cluster, provenance, and seed family identity, respectively; *β*_4_*VOC_Class*_*l*_ is the fixed effect of VOC class; *β*_5_(*Cluster*_*i*_ × *VOC_Class*_*l*_), *β*_6_(*Provenance*_*j*(*i*)_ × *VOC_Class*_*l*_), and *β*_7_(*Seed Family*_*k*(*j*(*i*))_ × *VOC_Class*_*l*_) represent the interactions between each hierarchical level of genetic background and VOC class; *μ*_*m*_ is the random intercept associated with Feature_ID *m*, assumed to follow a normal distribution 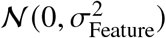; and *ε*_*ijklm*_ is the residual error term, assumed to be normally distributed *N* (0, *σ*^2^). Because of the nested relationship among cluster, provenance and seed family, type-I analysis of variance (ANOVA) was used to assess the significance of the fixed effects. Note that this is equivalent to specifying random terms as class/provenance/seed family in mixed-model notations Schmid et al., 2017. The advantage of using the fixed-terms approach is that it better allows to look at effects of individual clusters, provenances, and seed families (appropriate in the case of small numbers of groups) as well as their contributions to overall explained variance (in terms of increases in multiple R2 as measured by sum-of-squares proportions in the statistical model).

To further examine the effect of genetic background on each detected feature, we estimated the variation of VOCs explained by cluster, provenance, and seed family using the percentage of sum of squares (ss% as effect-size measure) from separated linear models fitted for each feature:

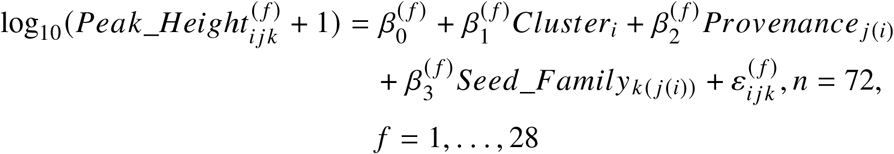

where 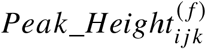 indicates the peak height of feature *f* measured in seed family *k* nested within provenance *j* and cluster *i*. Separate linear models were fitted for each feature (*f* = 1, …, 37). 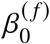 is the intercept for feature *f*, 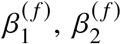, and 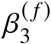 represent the fixed effects of cluster, provenance (nested within cluster), and seed family (nested within provenance), respectively, and 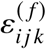 is the residual error term. We used type-I analysis of variance (ANOVA) to assess the significance of cluster, provenance within cluster, and seed family within provenance. The “emmeans” R package ((version 1.11.2-8, Lenth, 2025)) was used to compare the mean value of VOC peak height among the clusters across the VOC classes.

Furthermore, for features whose peak heights were significantly affected by genetic cluster or provenance, the data were analyzed in two ways: (1) using the peak height value (peak height > 0), and (2) coding detected samples as “1” and non-detected samples as “0”. Linear models were fitted to dataset (1), and generalized linear models (GLMs) were applied to dataset (2). Multiple group comparisons were conducted using the *emmeans* R package (version 1.11.2-8, Lenth, 2025) by Tukey’s HSD test (*α* = 0.05). The “cld” function from Multicomp R package (version 1.4.28, Hothorn et al., 2008) was used to indicate statistic significance. For dataset (1), the linear model was:

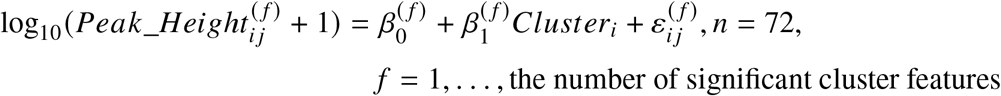

where 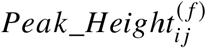 indicates the peak height (> 0) of feature *f* measured in cluster *i*. Separate linear models were fitted for each feature significant for cluster or provenance. 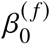 is the intercept for feature *f*, 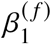 represents the fixed effect of cluster and 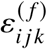 is the residual error term. When testing provenance instead of cluster, the term *Cluster*_*i*_ was replaced by *Provenance*_*i*_ in the model. For dataset (2), the generalized linear model is:

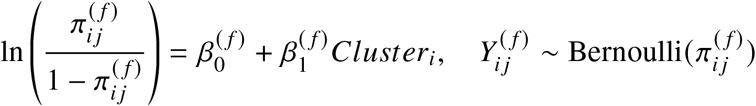

where 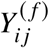 in 0, 1 represents the presence or absence of feature *f* for observation *j* in cluster *i*, 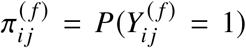 is the probability of feature occurrence, and *π*/(1 − *π*)) is the logit link function, mapping probabilities to the real line (−∞, ∞). When testing provenance instead of cluster, the term *Cluster*_*i*_ was replaced by *Provenance*_*i*_ in the model. Note that when provenance is fitted without cluster, it includes variation between provenances between clusters, not only between provenances within clusters.

##### Drought effects on VOC variation

To examine how European beech VOC profiles were distributed across three sample periods, we performed a principal component analysis (PCA) for each sampling period using the separately filtered data. The “prcomp” function in R was used for the PCA analysis after standardizing the variables to mean zero and unit standard deviation.

To statistically detect how drought affects European beech VOC profiles (H2), we analyzed the data in two complementary ways. First, we conducted a horizontal comparison of volatile profiles across periods, contrasting the drought and rewatering periods with the pre-drought period (Fig. 1c). Second, we analyzed drought and rewater periods separately using the individually filtered datasets.

For the horizontal comparison, we fitted an overall linear mixed-effects model (LMM) including all features for the drought and rewatering periods, respectively. The pre-drought peak height (log_10_-transformed) was included as a covariate at the beginning of the model. This approach allowed the model to first account for variation explained by baseline (pre-drought) peak height and then test for differences between the drought and rewatering periods relative to the pre-drought period. In other words, treatment effects were evaluated based on the remaining variation after controlling for pre-drought levels.

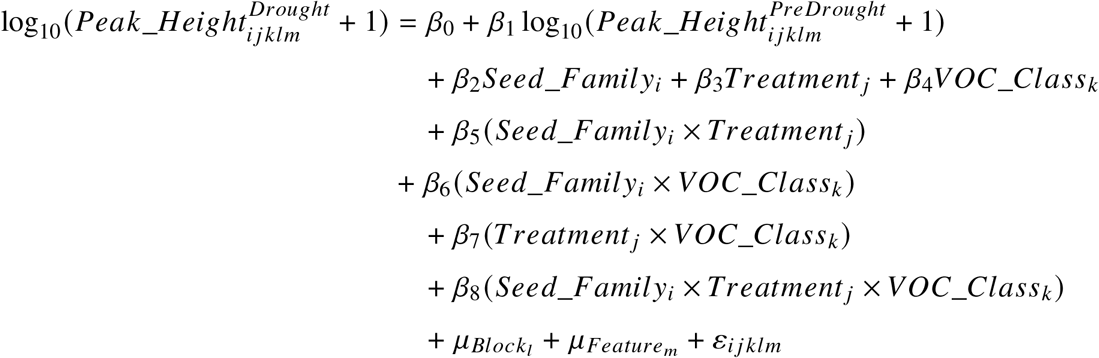

where log_10_(*Peak_Height*^*Drought*^*ijklm* + 1) is the log 10–transformed peak height (with a +1 offset) of feature *m* measured during the drought period, and log_10_(*Peak_Height*^*PreDrought*^*i j klm*+ 1) is the corresponding pre-drought (baseline) peak height included as a covariate; *β*_0_ is the intercept; *β*_1_ represents the effect of pre-drought peak height (baseline adjustment); *β*_2_*Seed_Family*_*i*_, *β*_3_*Treatment* _*j*_, and *β*_4_*VOC*_*C*_*lass*_*k*_ denote the fixed effects of seed family identity, treatment, and VOC class, respectively; *β*_5_(*Seed_Family*_*i*_ × *Treatment* _*j*_), *β*_6_(*Seed_Family*_*i*_ × *VOC_Class*_*k*_), and *β*_7_(*Treatment* _*j*_ ×*VOC_Class*_*k*_) represent the two-way interactions among seed family, treatment, and VOC class, while *β*_8_(*Seed_Family*_*i*_ ×*Treatment* _*j*_ ×*VOC_Class*_*k*_) denotes their three-way interaction; 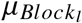 is the random intercept associated with block *l*, assumed to follow a normal distribution 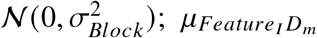 is the random intercept associated with feature identity *m*, assumed to follow 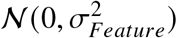; and *ε*_*ijklm*_ is the residual error term, assumed to be normally distributed *N* (0, *σ*^2^). When testing the rewatering period in comparison to the pre-drought period (instead of the drought period), the dependent variable was changed to 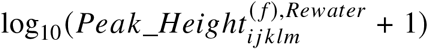, while the model structure and covariate adjustment remained the same. Type-I analysis of variance (ANOVA) was used to ensure that variation explained by the pre-drought covariate was fitted first in the sequential model. The “emmeans” R package (version 1.11.2-8, Lenth, 2025) was used to compare the mean value of VOC peak height between drought and control treatments across the VOC classes. Note that here and below when seed family is fitted without genetic cluster and provenance, it includes those. That is, when only seed family is fitted for the genetic background, it combines variation between seed families between clusters with variation between seed families between provenances within clusters and with variation between seed families within provenances.

To further compare the height peak change from pre-drought period to drought and re-water for each VOC feature, we applied the same test to each feature separately:

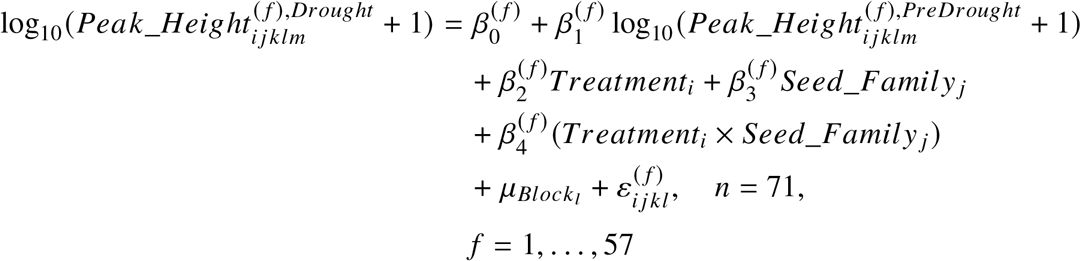

where 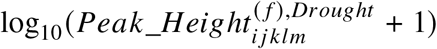 is the log_10_–transformed peak height of feature *f* measured during the drought period, and 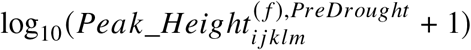 is the corresponding pre-drought peak height included as a covariate; 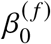 is the feature-specific intercept; 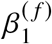 represents the effect of pre-drought peak height (baseline adjustment); 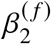 *Treatment*_*i*_ and 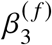 *Seed_Family* _*j*_ denote the fixed effects of Treatment and Seed_Family for feature *f*, respectively, and 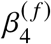 (*Treatment*_*i*_ × *Seed_Family*_*j*_) represents their interaction; 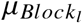 is the random intercept associated with block *l*, assumed to follow a normal distribution 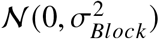, and 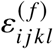 is the residual error term for feature *f*, assumed to follow a normal distribution *N* (0, *σ*^2^). Each feature was analyzed separately (*f* = 1, …, *F, n* = 71 per feature). One tree sample was lost during the drought period due to logistical reasons, resulting in a total of 71 tree samples during this period. When testing the re-water period in comparison to the pre-drought period (instead of the drought period), the dependent variable was changed to 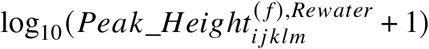 and *n* = 72, while the model structure and covariate adjustment remained the same. Type-I analysis of variance (ANOVA) was used to ensure that variation explained by the pre-drought covariate was fitted first in the sequential model.

For the individual period analyses, similar to the analyses for the pre-drought period, we used VOC data from 72 trees collected during drought treatment and re-water periods to fit an overall linear mixed-effects model, respectively. In the overall model, *log*_10_-transformed peak height was specified as the dependent variable, while treatment, seed family (as the overall genetic effect), as well as VOC class and their interactions with VOC class, were included as fixed effects. Block and Feature ID were included as random factors:

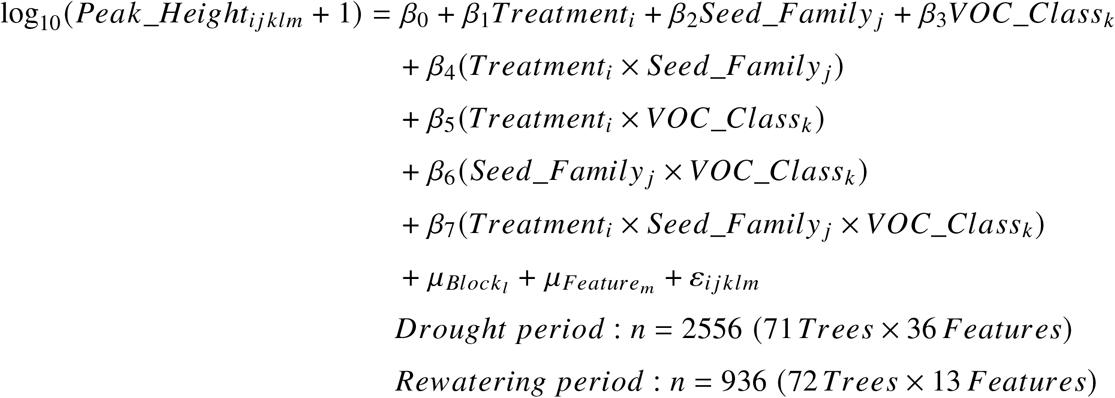

where log_10_(*Peak_Height*_*ijklm*_ + 1) is the log_10_–transformed peak height of feature *m* belonging to VOC class *k*, measured under treatment *i*, in seed family *j*, and block *l*; *β*_0_ is the intercept; *β*_1_*Treatment*_*i*_, *β*_2_*Seed_Family* _*j*_, and *β*_3_*VOC_Class*_*k*_ represent the fixed effects of drought treatment, seed family identity, and VOC class, respectively; *β*_4_(*Treatment*_*i*_ × *Seed_Family* _*j*_), *β*_5_(*Treatment*_*i*_ × *VOC_Class*_*k*_), and *β*_6_(*Seed*_*F*_*amily*_*j*_ × *VOC_Class*_*k*_) denote the two-way interactions among treatment, seed family, and VOC class, while *β*_7_(*Treatment*_*i*_×*Seed_Family* _*j*_ × *VOC_Class*_*k*_) represents their three-way interaction; 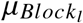 is the random intercept associated with block *l*, assumed to follow a normal distribution 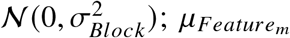 is the random intercept associated with feature identity *m*, assumed to follow 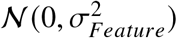; and *ε*_*ijklm*_ is the residual error term, assumed to follow *N* (0, *σ*^2^). Fixed effects were evaluated using Type-I ANOVA.

To further examine the genetic background effect on each detected features, we estimated the variation of VOCs explained by treatment, seed family, and their iteraction using the percentage of sum of squares (ss%) from separated linear models fitted for each feature in the drought period and the re-water period, respectively:

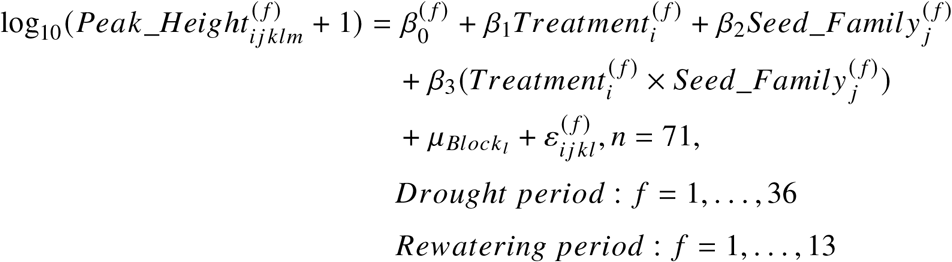

where log_10_(*Peak_Height*_*ijklm*_ + 1) is the log_10_–transformed peak height measured under treatment *i*, in seed family *j*, and block *l*; *β*_0_ is the intercept; *β*_1_*Treatment*_*i*_ and *β*_2_*Seed*_*F*_ *amil y* _*j*_ represent the fixed effects of drought treatment and seed family identity, respectively; and *β*_3_(*Treatment*_*i*_ × *Seed_Family* _*j*_) denotes their interaction; 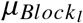 is the random intercept associated with block *l*, assumed to follow a normal distribution 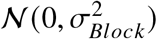, and *ε*_*ijkl*_ is the residual error term, assumed to be normally distributed *N* (0, *σ*^2^). Here, we used the aov function in R to fit the linear mixed-effects models (LMMs), which allowed greater flexibility in model specification and enabled us to explicitly test the treatment effect against block (Schmid et al., 2017).

Furthermore, for features whose peak heights were significantly affected by treatment, the data were analyzed in two ways: (1) using only detected samples (peak height > 0), and (2) coding detected samples as “1” and non-detected samples as “0”. Linear models were fitted to dataset (1), and generalized linear models (GLMs) were applied to dataset (2). Multiple group comparisons were conducted using the *emmeans* R (version 1.11.2-8, Lenth, 2025) package by Tukey’s HSD test (*α* = 0.05). The “cld” funtcion from the Multicomp R package was used to display letters for the statistical significance (cite). For dataset (1), the linear model is:

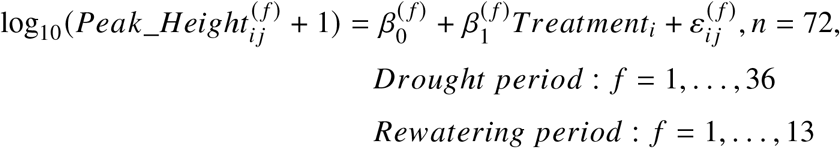

where 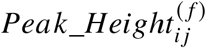 indicates the peak height (> 0)of feature *f* measured in cluster *i*. Separate linear models were fitted for each feature significant for treatment, where 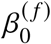 is the intercept for feature *f*, 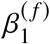 represent the fixed effects of treatment, and 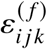 is the residual error term. For dataset (2), the generalized linear model is:

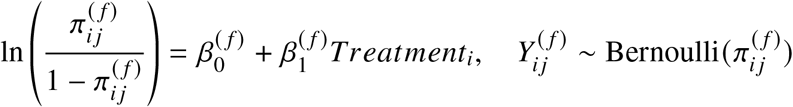

where 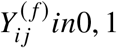 represents the presence or absence of feature *f* for observation *j* in treatment *i*; 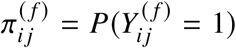 is the probability of feature occurrence; and *π*/(1 − *π*)) is the logit link function, which transforms probabilities bounded between 0 and 1 to an unbounded scale, enabling linear modeling of presence/absence data.

## Results

### Drought treatment was effective

Before the drought treatment, soil moisture and temperature were comparable between drought and control treatment (Supporting Information Fig. S2). The drought treatment resulted in reduced soil moisture in the drought compared to the control treatment, and soil temperature was also higher. By the time we conducted VOC sampling during the rewatering period, soil moisture and temperature were nearly identical again between the control and drought treatment (Supporting Information Fig. S2).

### Genetic background partially explains VOC variation, particularly in monoterpenes

Overall, genetic cluster and provenance within cluster significantly affected VOC peak height (Table 1). In addition, the interactions between cluster and VOC class, as well as between seed family (within provenance) and VOC class, were significant, genetic effects thus differed among VOC classes (Table 1).

**Table 1.** Results of the overall linear mixed-effects model testing the effects of genetic cluster, provenance, and seed family on VOC peak height. Cluster refers to the three genetic clusters, provenance to the specific site within a cluster, and seed family to the maternal origin within a site. VOC class refers to the classification of VOCs based on their biosynthetic pathways (eight classes were identified during the pre-drought period). Sum Sq and Mean Sq indicate sum square and mean square, respectively. NumDF and DenDF represent the numerator and denominator degrees of freedom, respectively. F value and P value indicate the F statistics and associated significance levels. Bold values in the ‘P value’ column indicate *P* < 0.05.

| Variable | Sum Sq | Mean Sq | NumDF | DenDF | F value | P value |
| --- | --- | --- | --- | --- | --- | --- |
| Cluster | 160.33 | 80.16 | 2 | 1900 | 24.69 | **<0.0001** |
| Provenance | 107.31 | 26.83 | 4 | 1900 | 8.26 | **<0.0001** |
| Seed family | 20.11 | 4.02 | 5 | 1900 | 1.24 | 0.2883 |
| VOC class | 5.03 | 0.72 | 7 | 20 | 0.22 | 0.9757 |
| Cluster:VOC class | 178.84 | 12.77 | 14 | 1900 | 3.93 | **<0.0001** |
| Provenance:VOC class | 109.00 | 3.89 | 28 | 1900 | 1.20 | 0.2179 |
| Seed family:VOC class | 218.69 | 6.25 | 35 | 1900 | 1.92 | **0.0010** |

Across individual feature-level models, genetic background explained 8.9% to 44.6% of the variation in VOC peak height (Fig. 2a). In total, 12 features were significantly affected by genetic cluster, nine by provenance within cluster, and nine by seed family within provenance. Genetic background significantly influenced seven monoterpenes, one oxidized terpenoid deriative, one GLV, two other fatty acid derivatives, and one unknown compound that could not be annotated (Fig. 2a).

**Figure 2.**
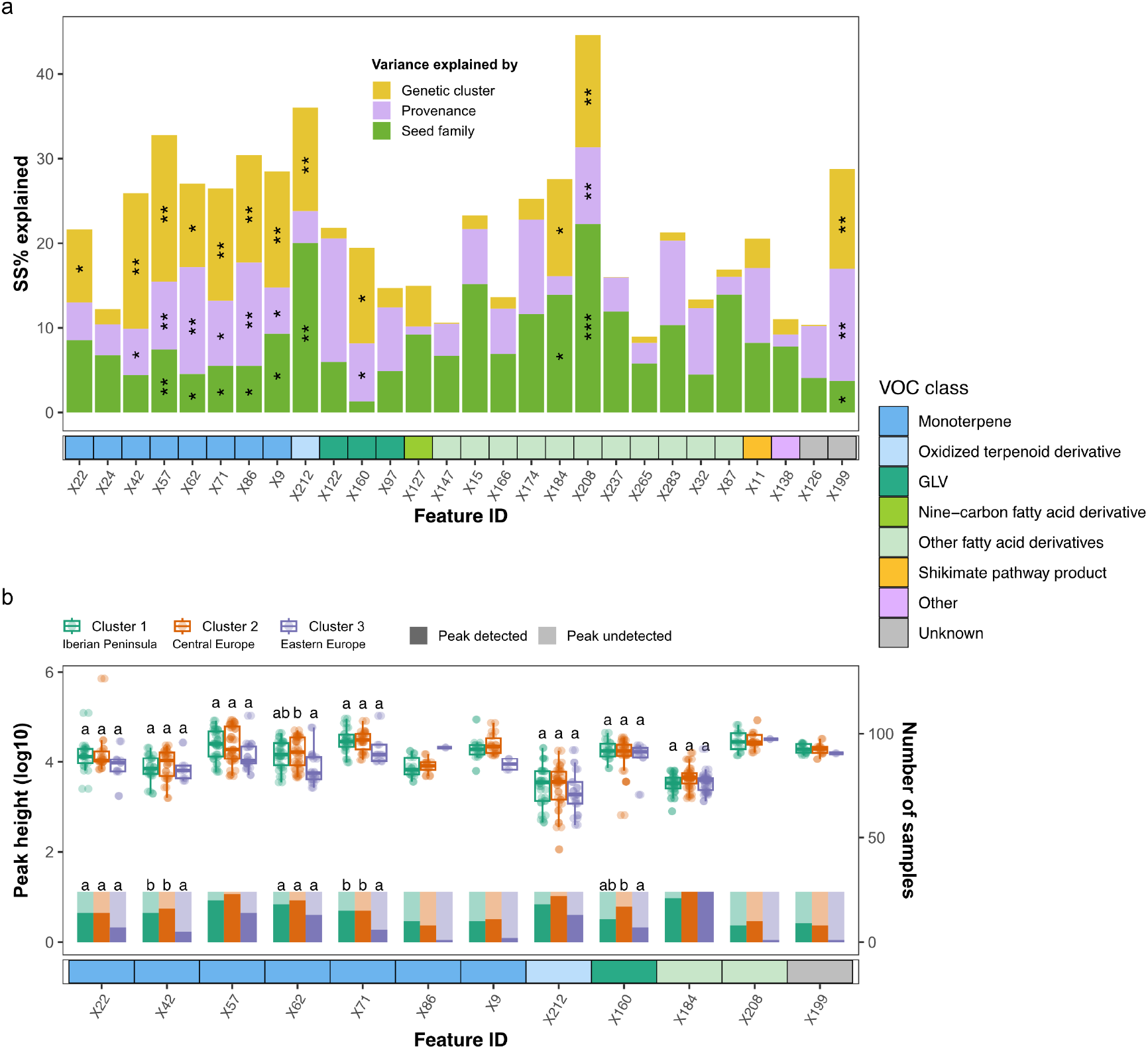
Effects of genetic background on VOC variation during the pre-drought period (n = 72 per feature). **(a)** Variation in VOC features explained by genetic cluster, provenance, and seed family, fitted sequentially in ANOVAs to calculate the percentage of sum of squares (SS%), representing the proportion of variance explained by each factor. Significance is indicated by asterisks on the bars (\**P* < 0.05; \*\**P* < 0.01; \*\*\**P* < 0.001 ; no asterisk indicates *P* ≥ 0.05). **(b)** Peak heights and detection frequencies of VOC features showing significant genetic cluster effects. Box plots show the distribution of peak heights for samples in which each feature was detected. The lower, middle, and upper hinges represent the first, median, and third quartiles (25th, 50th, and 75th percentiles), and whiskers extend to 1.5× the interquartile range. Bars below each box plot indicate the number of tree samples in which the feature was detected (dark color) or not detected (light color). Genetic clusters are distinguished by color. Different lowercase letters above the box plots and bars indicate significant differences among clusters based on Tukey’s HSD test (*α* = 0.05). Boxplots and bars without letters indicate that statistical comparisons were not performed because the feature was detected in fewer than three individuals in at least one genetic cluster (n < 3). Features’ classification is indicated by the colored bar below the x-axis.

Among the features significantly affected by genetic cluster, differences among the three clusters were observed in both detection frequency and peak height. Two monoterpenes (tentatively annotated as *β*-myrcene (x42) and *γ*-terpinene (x71) showed higher detection frequencies in the Iberian Peninsula and Central European than in the Eastern European cluster, while one green leaf volatile (tentatively annotated as 1-hexanol, 2-ethyl-(x160)) had a higher detection frequency in the Eastern European than in the Central European cluster (Fig. 2b). Among the samples in which it was detected, the mean peak height of one monoterpene (tentatively annotated as *β*-phellandrene (x62)) was lower in the Eastern European than in the Central European cluster (Fig. 2b). For provenance-level effects, peak height did not differ significantly among detected samples, and detection frequencies could not be statistically tested due to limited sample sizes (Supporting Information Fig. S3).

### Drought treatment modified VOC profiles

Principal component analysis (PCA) of VOC profiles across the three sampling periods showed clear differences in drought vs. control treatment separation over time (Fig. 3). During the pre-drought period, trees assigned to the drought and control treatments largely overlapped along both PC1 and PC2 axes (Fig. 3a). During the drought period, the two treatments exhibited substantially different distributions along both axes, with beech saplings in the drought treatment showing reduced variation. During rewatering, the drought and control treatments overlapped more than during the drought period, but remained separated compared with the pre-drought period (Fig. 3).

**Figure 3.**
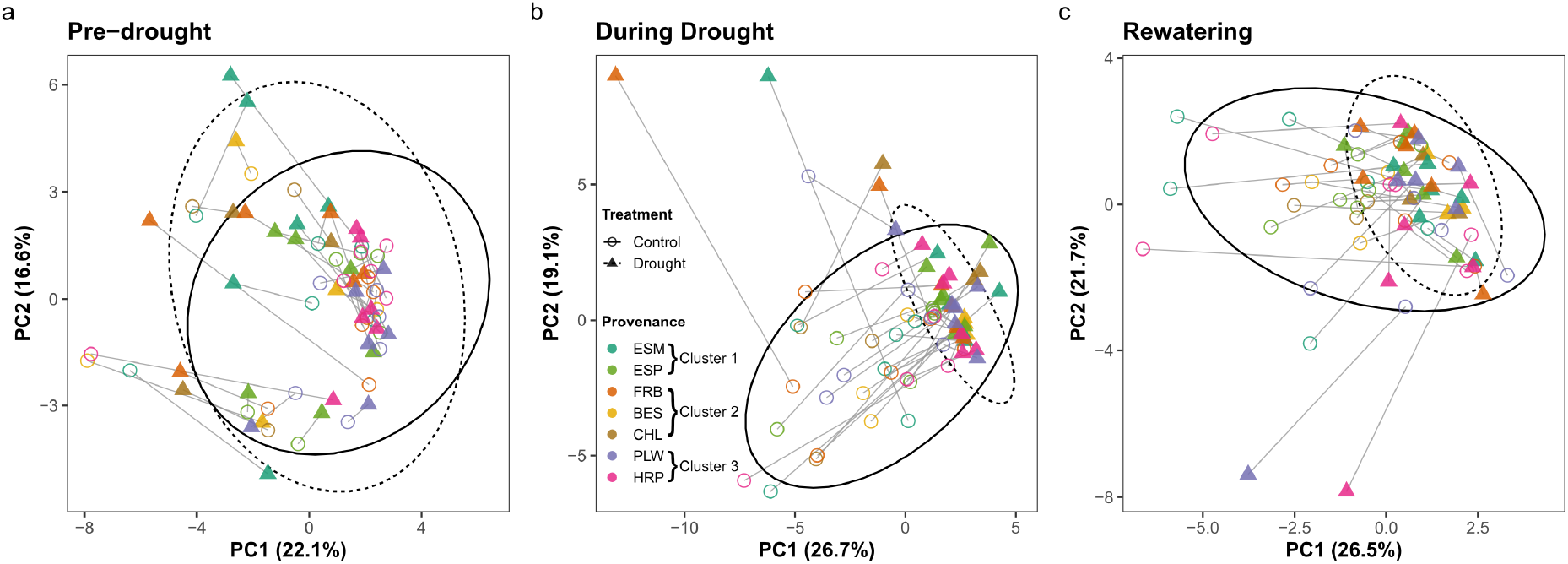
Principal component analysis (PCA) of detected VOC features across three sampling periods. **(a)** Pre-drought, **(b)** during drought, and **(c)** rewatering. PCA was performed separately for each sampling period. Control saplings are shown as open circles and drought-treated saplings as filled triangles. Saplings occupying the same block position and belonging to the same seed family but assigned to drought vs. control treatments are connected by grey lines. Ellipses represent 95% confidence intervals for each treatment group (dashed lines: drought; solid lines: control). Provenances are distinguished by color and arranged by the genetic clusters.

Overall, VOC peak heights during the drought period corresponded significantly to the VOC peak heights measured in the pre-drought period (Table 2). After controlling for this baseline effect, drought treatment significantly affected the change in VOC peak height, and the magnitude of this effect varied among VOC classes (Table 2).

**Table 2.** Results of the overall linear mixed-effects model assessing the effects of pre-drought period VOC peak height, seed family, drought treatment, VOC class, and their interactions on (a) VOC peak height during drought and (b) rewatering period. VOC peak height from the previous period was included as a covariate to account for baseline differences. Only seed family was included in the model to simplify the analysis; in this case seed-family varition includes variation among clusters and provenances. Sum Sq and Mean Sq indicate sum square and mean square, respectively. NumDF and DenDF represent the numerator and denominator degrees of freedom, respectively. F value and P value indicate the F statistics and associated significance levels. Bold values in the ‘P value’ column indicate *P* < 0.05.

| Variable | Sum Sq | Mean Sq | NumDF | DenDF | F value | P value |
| --- | --- | --- | --- | --- | --- | --- |
| **a) During drought period** |  |  |  |  |  |  |
| PeakHeight pre-drought | 40.90 | 40.90 | 1 | 119.38 | 13.70 | **0.0003** |
| Seed family | 64.54 | 5.87 | 11 | 3091.58 | 1.97 | **0.0278** |
| Drought treatment | 19.16 | 19.16 | 1 | 5.89 | 6.42 | **0.0451** |
| VOC class | 47.75 | 6.82 | 7 | 40.67 | 2.29 | **0.0464** |
| Seed family:Drought treatment | 95.20 | 8.65 | 11 | 3228.98 | 2.90 | **0.0008** |
| Seed family:VOC class | 405.46 | 5.27 | 77 | 3238.86 | 1.76 | **0.0001** |
| Drought treatment:VOC class | 974.12 | 139.16 | 7 | 3238.91 | 46.63 | **<0.0001** |
| Seed family:Drought treatment:VOC class | 271.21 | 3.52 | 77 | 3238.80 | 1.18 | 0.1369 |
| **b) Rewetting period** |  |  |  |  |  |  |
| PeakHeight pre-drought | 24.84 | 24.84 | 1 | 96.40 | 12.98 | **0.0005** |
| Seed family | 75.95 | 6.90 | 11 | 3109.99 | 3.61 | **<0.0001** |
| Drought treatment | 3.40 | 3.40 | 1 | 5.91 | 1.78 | 0.2314 |
| VOC class | 4.08 | 0.58 | 7 | 40.51 | 0.30 | 0.9478 |
| Seed family:Drought treatment | 43.90 | 3.99 | 11 | 3223.94 | 2.09 | **0.0183** |
| Seed family:VOC class | 263.04 | 3.42 | 77 | 3238.62 | 1.79 | **<0.0001** |
| Drought treatment:VOC class | 180.28 | 25.75 | 7 | 3238.66 | 13.46 | **<0.0001** |
| Seed family:Drought treatment:VOC class | 191.96 | 2.49 | 77 | 3238.57 | 1.30 | **0.0406** |

By analyzing individual VOC features during the drought period using the same baseline model, we found that eight monoterpenes, three oxidized terpenoid derivatives, three green leaf volatiles (GLVs), and three other fatty acid derivatives showed significant drought effects (Fig. 4). A separate analysis focusing only on the drought period using filtered data produced similar results, with drought treatment explaining up to 55.1% of the variation in peak height across features (Supporting Information Fig. S4a). All significant features were consistent between the two analyses, which explained 16.2% to 55.1% of VOC variation by drought treatment, except feature X86 (tentatively annotated as cyclohexene, 1-methyl-4-(1-methylethylidene)-), which was excluded from the drought period-only analysis due to insufficient detection during the drought period. Among these features, three monoterpenes and two other fatty acid derivatives showed lower detection frequencies, while three monoterpenes and two other fatty acid derivatives showed lower mean peak heights in the drought than in the control treatment. In contrast, two GLVs showed higher detection frequencies, and one GLV showed a higher mean peak height in the drought than in the control treatment (Supporting Information Fig. S4b).

**Figure 4.**
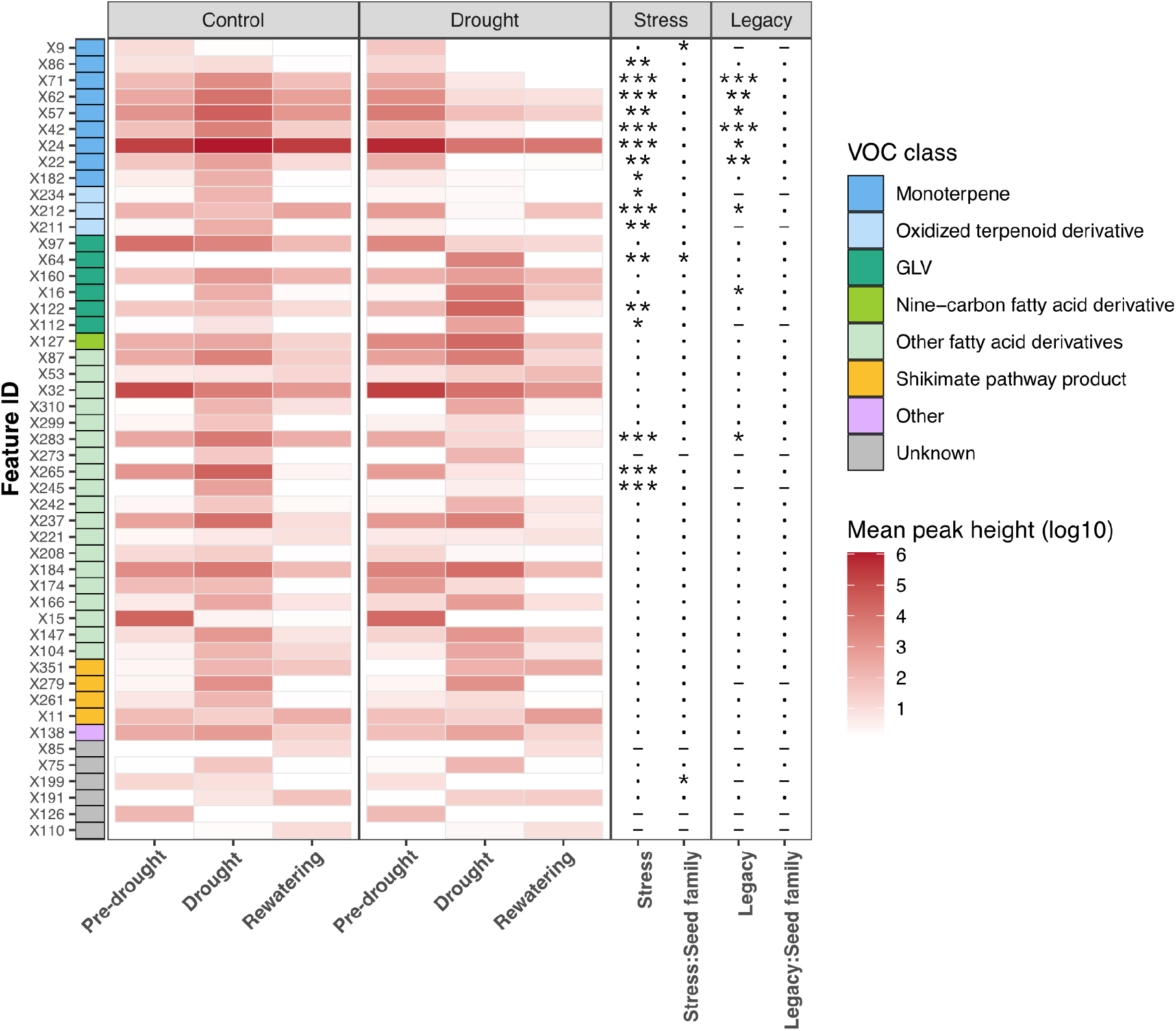
Mean VOC peak heights across three sampling periods, and the drought and legacy effects for each detected feature. Each row represents a VOC feature, and its classification is indicated by the colored bar on the y-axis. The log_10_-transformed mean peak height is shown by the darkness of the red color and is displayed separately for beech sapling of the drought and the control treatments across the pre-drought, drought, and rewatering periods. Drought and legacy effects, and their interactions with seed family, were tested as the treatment effect (drought *vs*. control) and its interaction with seed family on the change in feature peak height from pre-drought to drought and to rewatering, respectively. Significance is indicated by asterisks (\**P* < 0.05; \*\**P* < 0.01; \*\*\**P* < 0.001); a dot indicates *P ≥* 0.05; a dash indicates insufficient sample size for the corresponding statistical test.

During the drought period, seed family explained 2.9% to 32.6% of the variation of VOC feature height and was significant for one monoterpene, one oxidized terpenoid derivative, and two other fatty acid derivatives (Supporting Information Fig. S4a).

### Legacy effects of drought treatment on VOC profiles during rewatering

During the rewatering period, VOC peak heights were significantly influenced by the VOC peak heights measured in the pre-drought period (Table 2). Drought treatment alone did not significantly affect VOC peak height changes from pre-drought to the rewatering period; however, we detected a significant interaction between drought treatment and VOC class. This result shows a legacy effect of drought treatment on tree VOC profiles, with the strength of this legacy effect varying among VOC classes (Table 2).

During rewatering, six monoterpenes, one oxidized terpenoid derivative, and one other fatty acid derivative still showed significant legacy effects (Fig. 4). None of the VOC features showed a significant interaction between drought legacy effects and seed family (Fig. 4). We found 0.1% to 25.6% of the peak height variation for each feature still explained by drought treatment (Supporting Information Fig. S4a). Among these features, two monoterpenes and one other fatty acid derivatives showed lower detection frequencies in the drought treatment compared with the control treatment, while two monoterpenes showed lower mean peak heights under drought conditions (Supporting Information Fig. S5b).

During the rewatering period, seed family explained 8.2%-28.2% of the variation of VOC feature height and was only significant for two oxidized terpenoid derivative, and one other fatty acid derivative (Supporting Information Fig. S5a).

### Limited evidence for interaction effects between genetic background and drought treatment

During the drought period, seed family and drought interactions were significant in the overall mixed-effects model, indicating that seed families differed in their integrated VOC response to drought (Table 2). However, feature-level analyses only detected one monoterpene, one GLV, and one unknown feature with significant seed family-specific drought responses (Fig. 4). A similar pattern was observed during the rewatering period, where the overall seed family × drought interaction remained significant (Table 2), but no individual VOC feature showed a significant interaction effect (Fig. 4).

## Discussion

Using a common garden experiment with European beech saplings from diverse genetic backgrounds, we analyzed VOC emissions before drought, during drought, and after rewatering to disentangle genetic effects and drought-induced responses. Under non-drought conditions, genetic background across three hierarchical levels (genetic cluster, provenance within cluster, and seed family within provenances) explained a substantial proportion of VOC variation, indicating that baseline VOC profiles represent genetically determined chemical phenotypes. The strength of this genetic influence differed among VOC classes, with monoterpenes being particularly strongly differentiated between genetic background, suggesting that terpenoid biosynthesis is more tightly regulated by genetic factors than other compound groups. The drought treatment significantly altered VOC profiles, with the drought-treated saplings generally showing lower emissions of monoterpenes, oxidized terpenoid derivatives, and other fatty acid derivatives but higher emissions of GLVs. Furthermore, some drought-induced changes on monoterpenes persisted after rewatering, indicating a legacy effect of drought. This finding may reflect delayed physiological recovery or sustained metabolic adjustment reflecting drought stress. Although we detected significant seed family × drought treatment interactions when all VOC features were analyzed together, in separate analyses only three features showed significantly different drought responses among genetic backgrounds. Together, these results indicate that VOC phenotypes show relatively consistent responses to drought across genetic backgrounds and that therefore they may be suitable biomarkers to detect drought stress in European beech.

### Genetic background shapes VOC profiles

Our results show that genetic background explained a substantial proportion of variation in VOC peak heights. This finding supports our hypothesis and is consistent with previous studies demonstrating that plant VOC emissions can vary among genotypes and populations (reviewed by Niederbacher et al., 2015). Evidence from herbaceous and agricultural species shows that VOC production can be directly controlled by genetic background, including in *Arabidopsis thaliana* (Kang et al., 2025), *blueberry (Vaccinium spp*.) (Ferrão et al., 2020), tomato (*Solanum lycopersicum*) (Bauchet et al., 2017), and apple (*Malus domestica*) (Farneti et al., 2017). However, less is known about genetic differentiation in VOC emissions in long-lived broadleaved trees, with a few studies on cork oak (Staudt et al., 2025) and poplar (Eller et al., 2012). Our study extends this knowledge to European beech, an ecologically and economically important tree species in Europe.

Our results suggest that VOC profiles of European beech trees are structured by hierarchical genetic background, where genetic clusters and provenances reflect population differences shaped by both historical processes and local environmental adaptation, while seed families capture variation within populations, including potential maternal effects. This extends previous studies showing that traits such as leaf morphology and growth vary across multiple genetic levels, such as among genetic clusters (Kurath et al., 2025), among provenances (Müller and Finkeldey, 2016; Stojnić et al., 2016), and within provenances (Stojnić et al., 2016). The substantial within-provenance variation is further consistent with a previous study showed high within-population trait variation in European beech (Bresson et al., 2011).

Among VOC classes, monoterpenes were especially influenced by genetic background. Monoterpenes are a major component of VOC emissions in European beech (van Meeningen et al., 2016) and function in anti-herbivore defense, other stress responses, and pollinator attraction (Abbas et al., 2017; Baldwin, 2010; Zielińska-Błajet and Feder-Kubis, 2020). Their biosynthesis is regulated by terpene syntheses gene families (Abbas et al., 2017), and previous studies have demonstrated that monoterpene production is heritable and genetically regulated in grapes (*Vitis vinifrea*) (Zhang et al., 2025) and aromatic oil plants such as tea trees (*Melaleuca alternifolia*) (Voelker et al., 2023), sage (*Salvia officinalis, S. fruticosa and S. pomifera*) (Schmiderer et al., 2023) and mint (*Mentha suaveolens*) (Yang et al., 2024). Our results suggest that similar genetically structured regulation of monoterpene emissions may occur in European beech.

### VOCs can be used as biomarkers of drought stress

As we hypothesized, VOCs may be used as suitable biomarkers for detect drought stress in European beech trees. Drought stress significantly altered the VOC profiles of European beech trees, particularly affecting terpenoids (including monoterpenes and oxidized terpenoid derivatives) and fatty acid derivatives (including GLVs and other fatty acid derivatives). To date, only a few studies have investigated VOC responses to drought in major tree species in European forests, and these were largely limited to leaf-level measurements (Šimpraga et al., 2011; Bonn et al., 2019), narrow genetic backgrounds (Bonn et al., 2019; Fitzky et al., 2023), or only a single individual (Šimpraga et al., 2011). Our study improves current understanding by examining whole-plant VOC emissions across a wide range of genetic backgrounds.

In our study, drought-stressed beech saplings released less monoterpenes and oxidized terpenoid derivatives into the air than did unstressed saplings, and for monoterpenes this difference persisted to some degree after rewatering, indicating a legacy effect of drought stress. This is consistent with previous studies on older European beech saplings (8-10 years old) showing that monoterpene emissions decreased during drought stress (Bonn et al., 2019). Similar drought-induced reductions in monoterpene emissions have also been reported in other tree species, such as Scots pine (*Pinus sylvestris*) (Kreuzwieser et al., 2021) and oak (*Quercus ilex*) (Lavoir et al., 2009). Some studies, however, observed that European beech trees can also display a slight increase in monoterpene emissions under mild drought conditions (Wu et al., 2015) or at the beginning of a more severe drought (Šimpraga et al., 2011), likely because monoterpenes can function as protective antioxidants (Ormeño et al., 2007; Radwan et al., 2017). Under severe drought, a substantial reduction in photosynthetic carbon supply can constrain monoterpene biosynthesis (Brilli et al., 2007), leading to decreased monoterpene emissions (Loreto and Schnitzler, 2010). In our experiment, drought stress caused substantial leaf damage, including leaf yellowing and shedding, followed by an aseasonal flush of new leaves after rewatering in some trees (Fig.1c). As a result, photosynthetic capacity likely did not fully recover to pre-drought levels after rewatering, which may explain the observed legacy effect. The three affected oxidized terpenoid derivatives, tentatively annotated as *α*-thujenal (x211), bicyclo[3.1.0]hexan-2-one, 5-(1-methylethyl)-(x212) and *α*-terpineol (x234), originate from the oxidation of monoterpenes (Wise and Croteau, 1999); therefore, their decrease under drought may result from lower monoterpenes availability.

We also found that drought increased the emissions of GLVs but decreased three other fatty acid derivatives. The increase of GLVs under drought is consistent with previous studies on European Beech trees (Šimpraga et al., 2011; Fitzky et al., 2023). GLVs are fatty acid–derived volatiles produced from membrane lipids via the Lipoxygenase pathway (Feussner and Wasternack, 2002; Schuman, 2023) and can be released following membrane damage caused by drought (Capitani et al., 2009). In contrast, the reduction in other fatty acid derivatives (tentatively annotated as pentanoic acid (x245), 2h-pyran-2-one, tetrahydro-(x265) and cyclopentanone, 2-cyclopentylidene-(x283)) may reflect a shift in lipid metabolism, whereby fatty acid precursors are increasingly diverted to the production of epicuticular wax long-chain alkanes that serve as a hydrophobic barrier against water loss under drought stress. Increased wax accumulation under drought has previously been reported in European beech (Speckert et al., 2023), pine (*Pinus nigra*) (Kreyling et al., 2012), and grassland (*Holcus lanatus*) and heathland species (*Calluna vulgaris*) (Srivastava and Wiesenberg, 2018).

### VOC responses to drought are largely conserved across genetic backgrounds

Our results partially support our hypothesis that genetically different trees may differ in their VOC responses to drought (genotype × environment interaction = genetic variation in plasticity). We detected significant seed family × drought interactions both during the drought period and after rewatering in the overall model, while feature-level analyses identified significant interaction effects for only three features or none, respectively. Given that 57 VOC features were tested, approximately three significant results would be expected by chance alone at the 5% significance level. Thus, the feature-level results provide limited evidence for compound-specific genetic differences in drought responses. The significant overall interaction suggests that the interaction was driven by a few features or by small, coordinated shifts distributed across multiple compounds during drought and after rewatering. These findings are consistent with previous work showing that VOC emissions exhibit substantial phenotypic plasticity (environmental effects) but less genetic variation in phenotypic plasticity (genotype × environment interaction effects) in response to environmental factors, particularly drought (Niinemets, 2010). Although plants’ phenotypic plasticity is often expected to vary among genotypes as a result of local adaptation to climatic conditions (Nicotra et al., 2010; Aranda et al., 2018), our results are in line with previous studies in European beech finding that other functional traits, such as fine roots (Meier and Leuschner, 2008) and xylem traits (Unterholzner et al., 2025), are plastic to drought stress, with no significant difference in plasticity among provenances. Our findings imply that these plastic responses may be shaped by shared physiological mechanisms, such as reduced carbon assimilation and membrane stress (reviewed by Reddy et al., 2004), which commonly lead to similar metabolic adjustments across diverse genetic backgrounds. The conserved plasticity observed here indicates that VOC profiles may serve as indicators of drought stress that are largely independent of genetic background. Furthermore, the relatively low genetic variation in plasticity and absence of clear local specialization may also be related to the comparatively high within-vs. between-provenance genetic variation in European beech (Frank et al., 2017).

## Conclusions and outlook

We show that variation in VOC profiles under well-watered conditions reflects hierarchical genetic structure, while drought induces a conserved plastic response across genetic backgrounds, characterized by increased GLVs and decreased monoterpenes, oxidized terpenoid derivatives and other fatty acid derivatives. Persistently reduced monoterpenes after rewatering indicate legacy effects of drought. Because this study was conducted on potted saplings exposed to an experimental drought treatment, further research is needed to determine whether similar patterns occur in mature trees under natural field conditions. It is also important to note that the genetic clusters examined here represent phylogeographic lineages (Lazic et al., 2024; Stefanini et al., 2022). To better assess adaptive responses of European beech to drought, it would be necessary to separate geographic from environmental variation, e.g. by selecting populations on drought-vs. non drought-exposed sites within geographic regions and repeating this design in multiple geographic regions. Such experiments would ideally be conducted under more natural conditions to address the limitations inherent to pot experiments. Additionally, the VOC was collected after the saplings showed drought symptoms on leaves, further work could assess whether VOC responses occur before visible drought symptoms. Nevertheless, our findings demonstrate that VOC profiles reflect both genetic background and physiological responses to drought stress, highlighting their potential as sensitive and non-invasive biomarkers for detecting drought stress and improving predictions of forest responses to climate change.

## Acknowledgments

We thank Sergio E. Ramos for assistance with fieldwork and TD-GC-EI-MS measurements. We thank Christian Geckeler for providing the programmable pumps, and Alexander Steppke for designing and building the VOC sampling system, including the wooden frame and fan system, as well as improving the pump reprogramming methods. We also thank Luca Rohrbach and Sophie A. Schuman-Steppke for assistance in building the VOC sampling system. This work was supported by the NOMIS foundation grant Remotely Sensing Ecological Genomics to PI M. E. Schaepman, whom we thank for supporting this experiment and MCS as Co-Investigator. We further thank the financial support of the project The Significance of Aerosols and Volatile Organic Substances for the Incorporation of Plant Biomass into Soil through the Foundation for Research in Science and the Humanities at the University of Zurich to GLBW.

## Author contributions

Conceptualization: GLBW, MCS, SvM, TT. Data curation: TT. Formal analysis: LR, TT. Funding acquisition: MCS. Investigation: TT, TG, SvM. Methodology: GLBW, LR, MCS, SvM, TT. Project administration: MCS, SvM, GLBW. Resources: MCS, GLBW. Supervision: MCS, SvM. Validation: LR. Visualization: TT. Writing - original draft: TT. Writing - review and editing: All authors.

## Data availability

The data and R scripts used in this study have been deposited in Dryad, and the raw GC–MS data have been deposited in MetaboLights. The repositories are currently under embargo during peer review and will be made publicly available upon acceptance of the manuscript.

