## Supporting Information for "Drought alters volatile profiles in European beech saplings across genetically diverse backgrounds"

The following Supporting Information is available for this article:

**Table S1.** VOC groups and VOC classes found in this study.

**Table S2.** Paired comparisons of VOC peak heights between drought and control groups for each VOC class during the drought period.

**Table S3.** Tentative annotations, VOC groups and classes, spectral characteristics, and occurrence frequencies of detected VOC features.

**Table S4.** Major biosynthetic classes of plant volatiles, their characteristics, and reported functions.

**Fig. S1.** Overview of the genetic variation source, experimental design, and VOC sampling setup.

**Fig. S2.** Environmental moisture and temperature during the experiment.

**Fig. S3.** Peak heights of significant VOC features across the seven provenances.

**Fig. S4.** Drought treatment effects on VOC variation during the drought period.

**Fig. S5.** Drought treatment effects on VOC variation during the rewatering period.

### Supporting tables

**Table S1.** VOC group and VOC class that were found in this study

| Biosynthetic group | VOC class |
|---|---|
| Terpenoid | Monoterpene |
| | Oxidized terpenoid |
| Fatty acid derivative | GLV |
| | Nine-carbon fatty acid derivative |
| | Other fatty acid derivatives |
| Shikimate pathway product | Shikimate pathway product |
| Other | Other |
| Unknown | Unknown |
| Contaminant | Synthetic product |
| | Nylon decomposition product |
| | Column bleed |

**Table S2.** Paired comparison between the mean of VOC peak height between the drought and control group for each VOC class during the drought period. The bold numbers in 'P value' indicate P < 0.05.

| Contrast | VOC class | estimate | SE | Z ratio | P value |
|---|---|---|---|---|---|
| Control - Drought | Monoterpene | 2.1953 | 0.2488 | 8.8226 | <**0.0001** |
| Control - Drought | Oxidized terpenoid derivative | 1.9432 | 0.3154 | 6.1609 | <**0.0001** |
| Control - Drought | GLV | -1.1447 | 0.2669 | -4.2892 | <**0.0001** |
| Control - Drought | Nine-carbon fatty acid derivative | -1.7293 | 0.4612 | -3.7497 | **0.0002** |
| Control - Drought | Other fatty acid derivatives | 0.4439 | 0.2279 | 1.9480 | 0.0514 |
| Control - Drought | Shikimate pathway product | 0.2056 | 0.2921 | 0.7039 | 0.4815 |
| Control - Drought | Other | 0.2373 | 0.4608 | 0.5149 | 0.6066 |
| Control - Drought | Unknown | -0.0425 | 0.2669 | -0.1592 | 0.8735 |

**Table S3.** Tentative annotations, volatile groups, volatile class, spectral characteristics(m/z and retention time (RT)) and the number of samples (No. sample) of VOC features detected after filtering rare features.

| Feature ID | Tentative annotation | VOC group | VOC class | m/z | RT | No. sample |
|---|---|---|---|---|---|---|
| X9 | $\gamma$-Pinene | Terpenoid | Monoterpene | 93.037 | 4.130 | 25 |
| X11 | Toluene | Shikimate pathway product | Shikimate pathway product | 91.050 | 4.297 | 225 |
| X15 | Acetic acid, butyl ester | Fatty acid derivative | Other fatty acid derivatives | 43.046 | 4.819 | 90 |
| X16 | Hexanal | Fatty acid derivative | GLV | 44.055 | 4.906 | 247 |
| X22 | Bicyclo[3.1.0]hexane, 4-methylene-1- | Terpenoid | Monoterpene | 93.060 | 5.296 | 76 |
| X24 | Bicyclo[3.1.0]hexane, 4-methylene-1- | Terpenoid | Monoterpene | 93.074 | 5.506 | 201 |
| X32 | 3-Hexenal | Fatty acid derivative | Other fatty acid derivatives | 56.084 | 5.806 | 216 |
| X42 | $\beta$-Myrcene | Terpenoid | Monoterpene | 91.057 | 6.193 | 87 |
| X53 | Heptanal | Fatty acid derivative | Other fatty acid derivatives | 70.071 | 6.435 | 105 |
| X57 | D-Limonene | Terpenoid | Monoterpene | 68.063 | 6.686 | 147 |
| X62 | $\beta$-Phellandrene | Terpenoid | Monoterpene | 77.030 | 6.821 | 127 |
| X64 | 2-Hexenal, | Fatty acid derivative | GLV | 55.076 | 6.906 | 30 |
| X71 | $\alpha$-Terpinene | Terpenoid | Monoterpene | 93.070 | 7.401 | 83 |
| X75 | unknown | unknown | unknown | 104.092 | 7.497 | 74 |
| X85 | unknown | unknown | unknown | 55.051 | 7.927 | 31 |
| X86 | Cyclohexene, 1-methyl-4- | Terpenoid | Monoterpene | 93.061 | 7.961 | 29 |

| | | | | | | |
|---|---|---|---|---|---|---|
| X87 | Octanal | Fatty acid derivative | Other fatty acid derivatives | 68.064 | 8.003 | 312 |
| X97 | 4-Hexen-1-ol, acetate | Fatty acid derivative | GLV | 43.058 | 8.420 | 123 |
| X104 | 5-Hepten-2-one, 6-methyl- | Fatty acid derivative | Other fatty acid derivatives | 108.100 | 8.673 | 130 |
| X110 | unknown | unknown | unknown | 67.068 | 8.835 | 63 |
| X112 | 1-Hexanol | Fatty acid derivative | GLV | 56.099 | 8.880 | 29 |
| X122 | 3-Hexen-1-ol, | Fatty acid derivative | GLV | 41.090 | 9.309 | 94 |
| X126 | unknown | unknown | unknown | 58.050 | 9.449 | 38 |
| X127 | Nonanal | Fatty acid derivative | Nine-carbon fatty acid derivative | 57.081 | 9.497 | 349 |
| X138 | Acetic acid | Other | Other | 43.058 | 10.089 | 215 |
| X147 | 1-Tetradecene | Fatty acid derivative | Other fatty acid derivatives | 69.072 | 10.288 | 179 |
| X159 | 3-Bromooctane | Contaminant | Synthetic product | 57.091 | 10.687 | 72 |
| X160 | 1-Hexanol, 2-ethyl- | Fatty acid derivative | GLV | 57.092 | 10.758 | 310 |
| X166 | Decanal | Fatty acid derivative | Other fatty acid derivatives | 57.080 | 10.912 | 342 |
| X174 | Propanoic acid | Fatty acid derivative | Other fatty acid derivatives | 74.031 | 11.250 | 134 |
| X182 | Linalool | Terpenoid | Monoterpene | 69.043 | 11.511 | 32 |
| X184 | 1-Octanol | Fatty acid derivative | Other fatty acid derivatives | 56.089 | 11.614 | 243 |
| X191 | unknown | unknown | unknown | 81.082 | 11.815 | 99 |
| X199 | unknown | unknown | unknown | 56.065 | 12.209 | 25 |

| | | | | | | |
|---|---|---|---|---|---|---|
| X208 | Butyrolactone | Fatty acid derivative | Other fatty acid derivatives | 42.083 | 12.413 | 35 |
| X211 | $\gamma$-Thujenal | Terpenoid | Oxidized terpenoid derivative | 79.066 | 12.508 | 27 |
| X212 | Bicyclo[3.1.0]hexan-2-one, 5- | Terpenoid | Oxidized terpenoid derivative | 97.069 | 12.548 | 224 |
| X221 | 3-Octadecene, | Fatty acid derivative | Other fatty acid derivatives | 69.068 | 12.863 | 59 |
| X225 | 2-Pyrrolidinone, 1-methyl- | Contaminant | Synthetic product | 99.050 | 12.989 | 81 |
| X234 | $\gamma$-Terpineol | Terpenoid | Oxidized terpenoid derivative | 59.087 | 13.329 | 25 |
| X237 | Heptadecane | Fatty acid derivative | Other fatty acid derivatives | 57.092 | 13.507 | 129 |
| X242 | Dodecanal | Fatty acid derivative | Other fatty acid derivatives | 96.103 | 13.555 | 73 |
| X244 | [1,1’-Bicyclopentyl]-2-one | Contaminant | Synthetic product | 84.069 | 13.657 | 191 |
| X245 | Pentanoic acid | Fatty acid derivative | Other fatty acid derivatives | 60.048 | 13.731 | 26 |
| X261 | Methyl salicylate | Shikimate pathway product | Shikimate pathway product | 120.029 | 14.418 | 59 |
| X263 | 1,1,1,3,3,5,5,7,7,9,9,11,11,13,13,15,15,15-octadecamethyloctasiloxane | Contaminant | Column bleed | 73.041 | 14.643 | 43 |
| X265 | 2H-Pyran-2-one, tetrahydro- | Fatty acid derivative | Other fatty acid derivatives | 42.066 | 14.758 | 93 |
| X273 | Hexanoic acid | Fatty acid derivative | Other fatty acid derivatives | 60.050 | 15.438 | 71 |

| | | | | | | |
|---|---|---|---|---|---|---|
| X279 | Benzyl alcohol | Shikimate pathway product | Shikimate pathway product | 79.078 | 16.019 | 58 |
| X283 | Cyclopentanone, 2-cyclopentylidene- | Fatty acid derivative | Other fatty acid derivatives | 150.099 | 16.197 | 125 |
| X299 | 1-Dodecanol | Fatty acid derivative | Other fatty acid derivatives | 69.088 | 18.080 | 67 |
| X300 | 1,1,1,3,3,5,5,7,7,9,9,11,11,13,13,15,15,17,17,19,19,19-docosamethyldecasiloxane | Contaminant | Column bleed | 73.043 | 18.415 | 34 |
| X310 | Octanoic acid | Fatty acid derivative | Other fatty acid derivatives | 60.048 | 20.000 | 68 |
| X322 | Caprolactam | Contaminant | Nylon decomposition product | 55.053 | 22.463 | 223 |
| X325 | Caprolactam | Contaminant | Nylon decomposition product | 55.065 | 22.554 | 68 |
| X341 | Dibutyl adipate | Contaminant | Synthetic product | 129.064 | 25.095 | 207 |
| X351 | 2-Ethylhexyl salicylate | Shikimate pathway product | Shikimate pathway product | 120.026 | 25.780 | 103 |

**Table S4.** Major biosynthetic classes of plant volatiles, their characteristics, and reported functions, further expanded and updated from Schuman et al. (2016).

| Class | Compounds | Biosynthesis | Functions | Volatility (BP, 760 mm Hg)[b] | Known structures |
|---|---|---|---|---|---|
| Fatty acid derivatives | Jasmonates | From 16:3 and 18:3 fatty acids dioxygenated at C13 by 13-LOX (Wasternack, 2007) | Floral scent (Demole et al., 1962) and volatile forms of plant hormones (Karban et al., 2000; Preston et al., 2001; Kessler et al., 2006; Birkett et al., 2000) | (*Z*)-jasmone, 291°C; methyl jasmonate, 303°C | Four stereo-isomers |
| | Green leaf volatiles (GLVs: six-carbon volatile aldehydes, alcohols, and esters) | From cleavage of 13-LOX products by HPL to yield hexenal (from 18:2) or (*Z*)-3-hexenal (from 18:3 fatty acids), which may be converted to alcohols by ADH, and further esterified (Matsui, 2006) | Typical damaged leaf or "cut grass" smell (Hatanaka et al., 1987), also emitted from other organs (Dudareva et al., 2006); antimicrobial or antifungal (Deng et al., 1993; Shiojiri et al., 2006); may stimulate animal consumption as flavor components (Halitschke et al., 2004); part of direct (Vancanneyt et al., 2001) and indirect anti-herbivore defense (Shiojiri et al., 2006), and may prime or elicit defense within (Frost et al., 2008) and between plants (Paschold et al., 2006; Baldwin et al., 2006; Sugimoto et al., 2014) | (*Z*)-3-hexenal, 122.7°C; (*Z*)-3-hexenol, 156.5°C; (*Z*)-3-hexenyl acetate, 191°C | Tens: four aldehydes (hexanal, (*Z*)-3-hexenal, (*E*)-2-hexenal, (*E*)-3-hexenal), thus four alcohols, each potentially with esters including acetates, propionates, butyrates, isobutyrates, valerates, isovalerates, benzoates, salicylates |
| | Nine-carbon volatile aldehydes, alcohols and esters | From 9-LOX products of 18:2 and 18:3 fatty acids, HPL and ADH. Some HPLs cleave only 9- or 13-hydroperoxides, while others cleave both. Products from 18:2 have one double bond, and from 18:3 have two (De Domenico et al., 2007). | Fruit odor and flavor components (Vancanneyt et al., 2001), antifungals (Matsui, 2006); possibly involved in almond seed development (Mita et al., 2005) | (*E*,*E*)-3,6-nonadienal, 202°C; (*E*,*E*)-3,6-nonadienol, 215°C; (*E*,*E*)-3,6-nonadienyl acetate, 247°C | Tens: five aldehydes and thus five alcohols, which can be esterified; acetate esters most frequently reported |
| | Pentyl leaf volatiles (PLVs: five-carbon ketones, aldehydes, alcohols, and esters) (Fall et al., 2001; Huang et al., 2025) | A C5-C13 oxidative cleavage of 18:2 and 18:3 fatty acids by lipoxygenase can produce a pentane (18:2) or pentene (18:3) radical that reacts to form alcohols, aldehydes, esters, and ketones (Fall et al., 2001; Gorman et al., 2021). Note that pentanal may also be derived from pyruvate (Wang et al., 2019). | As for GLVs, PLV emission is associated with tissue damage (Fall et al., 2001), light-dark transitions (Jardine et al., 2012), and defenses against insects and pathogens (Huang et al., 2025). PLVs are expected to be more abundant than GLVs under low-oxygen conditions (Fall et al., 2001). | pentanal, 104°C; 1-penten-3-one, 150°C; (*Z*)-2-pentenyl acetate, 151°C | Tens: three aldehydes and one ketone and thus four alcohols, which can be esterified; acetate esters most frequently reported |

| Class | Compounds | Biosynthesis | Functions | Volatility (BP)[b] | Known structures |
|---|---|---|---|---|---|
| | Other fatty acid derivatives (ketones, shorter-chain fatty acids, aldehydes, alcohols, esters, alkanes, alkenes) | Oxidative cleavage and decarboxylation of various fatty acids also results in shorter-chain or medium-chain volatiles with aldehyde and ketone moieties and derivatives of these (Howe and Schilmiller, 2002; Pichersky et al., 2006; D'Auria et al., 2002). Volatiles including alkanes, alkenes, aldehydes, ketones, and lactones are also derived from cuticular waxes (Chen et al., 2023). Note that volatile short-chain fatty acids are generally the product of digestion or fermentation and not generally reported to be emitted by plants; see *e.g.* (Weinhold and Baldwin, 2011). | Many shorter-chain aldehydes and ketones serve as precursors for the biosynthesis of other volatiles (Pichersky et al., 2006) while methylketones are well described as defensive compounds from plant glandular trichomes (Fridman et al., 2005). | 2-heptanone, 150°C; 2-undecanone, 232°C; 2-pentadecanone, 293°C | Hundreds to thousands of structures |
| Terpenoids | Canonical terpene hydrocarbons: hemiterpenes (C5), monoterpenes (C10), sesquiterpenes (C15), and some diterpenes (C20) (Yáñez-Serrano et al., 2018) | From 5-carbon precursors IPP and DMAPP via one of two pathways in plants: the MEP pathway in plastids or the MVA pathway in the cytosol. Generally, hemiterpenes and monoterpenes are synthesized in plastids and sesquiterpenes in the cytosol; some sesquiterpenes may be synthesized in mitochondria from cytosolic substrate (Rodríguez-Concepción, 2006; Kappers et al., 2005); production is usually light-dependent (Lerdau and Gray, 2003). | Odors from green tissue, flowers and fruits, and roots; many reported to be allelopathic (Mizutani, 1999), antimicrobial or antifungal (Cowan, 1999; Khosla and Keasling, 2003), function as direct (Brattsten, 1983) or indirect (Degenhardt et al., 2003) antiherbivore defenses, attract pollinators (Schiestl, 2010), and be involved in defense elicitation and priming (Arimura et al., 2000). Most react with atmospheric ozone (Calogirou et al., 1999) and could be involved in plant oxidative stress responses (Vickers et al., 2009). | isoprene, 34°C; (*Z*)-($\beta$)-ocimene, 175°C; (*S*)-(-)-limonene, 177°C; (*E*)-($\beta$)-farnesene, 273°C; (-)-($\beta$)-caryophyllene, 263°C; kaurene, 347°C | Hemiterpenes: only isoprene; perhaps 1000 mono- and 5000 sesquiterpenes (Seigler, 2008; Gershenzon and Croteau, 1991), most mono- or polycyclic |
| | Norsesquiterpenes and norditerpenes (often referred to as homoterpenes) and apocarotenoids (C8–C18) | Norditerpenes and norsesquiterpenes are derived from diterpenes (C20) in plastids or sesquiterpenes (C15) in the cytosol by oxidation, possibly catalyzed by Cyp450 (Herde et al., 2008; Chappell and Coates, 2010; Dudareva et al., 2006); apocarotenoids are cleaved from carotenoids in plastids by CCO (Auldridge et al., 2006; Walter et al., 2010). | (*E*,*E*)-TMTT and (*E*)-DMNT are commonly reported to mediate indirect defense of leaves (Dudareva et al., 2006). Apocarotenoids are flavor and odor components of fruits, flowers, and green tissue, reported as both attractants and repellents of pollinators and predators, and associated with fruit ripening (Camara and Bouvier, 2004; Bouvier et al., 2005); some are anti-fungal (Maffei, 2010). | (*E*)-DMNT, 196°C; (*E*,*E*)-TMTT, 293°C; $\beta$-ionone, 282°C | Only (*E*)-DMNT and (*E*,*E*)-TMTT are widely reported from plants, although a handful of other homoterpenes are reported from insects (Wegener and Schulz, 2002) |

| Class | Compounds | Biosynthesis | Functions | Volatility (BP)[b] | Known structures |
|---|---|---|---|---|---|
| | Oxidized terpenoids and derivatives | Derived from terpenes by oxidation, *e.g.* by Cyp450. Products may be further oxidized, esterified, or reduced; some TPS's synthesize oxidized terpenoids by incorporating $CO_2$ (Dudareva et al., 2006). | Odor components of fruits, flowers, green tissue, and roots (Dudareva et al., 2004) with similar ecological roles reported as for terpene hydrocarbons, but more often directly toxic (Khosla and Keasling, 2003); precursors of aerosols (Blande et al., 2014). | prenol, 142°C; linalool, 199°C; (*E*,*E*)-farnesol, 283°C | Similar as for terpene hydrocarbons |
| Shikimate pathway | Acid, aldehyde and alcohols derived from L-phenylalanine, indole (tryptophan precursor), and other derivatives of shikimate products | L-phenylalanine is converted to trans-cinnamic acid, with a C3 side chain, by PAL, and then to other phenylpropanoids by steps of monolignol biosynthesis, and the side chain may be enzymatically shortened to produce benzenoids, or other derivatives having a C2 side chain. Indole is a direct precursor of tryptophan (Dudareva et al., 2006). | Common in floral scents (Vogt, 2010) and source of capsaicinoids (pungence in pepper); methyl salicylate is a common herbivore-induced leaf volatile which attracts some predators and parasitoids (Van Poecke et al., 2001; Ament et al., 2004). | methyl salicylate, 222°C; indole, 253°C | ca. 20% of known plant volatiles (Qualley and Dudareva, 2008) |
| Other amino acid derivatives | Acids, aldehydes, alcohols, esters, nitrogen- and sulphur-containing volatiles from non-aromatic amino acids, ethylene (from methionine) (Dudareva et al., 2006; Maoz et al., 2022) and nitrous oxide (from arginine) | Amino acids are deaminated or transaminated to $\alpha$-keto acids, which are carboxylated and may be reduced, oxidated or esterified. Amino acids may also be precursors for acyl coA molecules used in esterification by alcohol acyltransferases (Dudareva et al., 2006). | Amino-acid derived esters are found in flowers and fruits (Maoz et al., 2022); branched-chain amino acid (Leu, Ile, Val) derivatives are common in fruit (Dudareva et al., 2006). Putrid sulphur-containing compounds, likely from methioinine (Maoz et al., 2022), may act as direct defences (Berkov et al., 2000). Ethylene and nitrous oxide are endogenous signals. | ethylene, -104°C; 2-methylbutanal, 94°C; 3-(methylthio)propyl acetate, 201°C | Unclear |

[a] In addition to these classes, methanol and acetic acid are produced abundantly from cell wall O-acetyl and methyl esterification processes during leaf development (Dewhirst et al., 2020), and other volatiles are produced from precursors of fatty acids and carbohydrates (Jardine et al., 2012). Within each class (e.g. fatty acid derivatives, terpenoids), the order of entries corresponds approximately to how well this class of compounds is characterized in the plant volatile literature; individual compounds may be well characterized or frequently reported even when the class as a whole is less well characterized.

[b]Royal Society of Chemistry ChemSpider: http://www.chemspider.com/, accessed 29 May 2022 or 24 Nov 2025 for PLVs and other fatty acid derivatives (note that BP for(*Z*)-2-pentenyl acetate is from the Good Scents Company, https://www.thegoodscentscompany.com/data/rw1537951.html, accessed 24 Nov 2025); BP: boiling point.

[tfn3]ADH, alcohol dehydrogenase; CCO, carotenoid cleavage oxidase; Cyp450, cytochrome P450; DMAPP, dimethylallyl pyrophosphate; (*E*)-DMNT, trans-4,8dimethyl-1,3,7nonatriene; IPP, isopentyl pyrophosphate; HPL, hydroperoxide lyase; LOX, lipoxygenase; MEP, 2-C-methyl-derythritol 4-phosphate; MVA, mevalonic acid; (*E*,*E*)-TMTT: *trans*, *trans*-4,8,12-trimethyltrideca-1,3,7,11-tetraene.

### Supporting figures

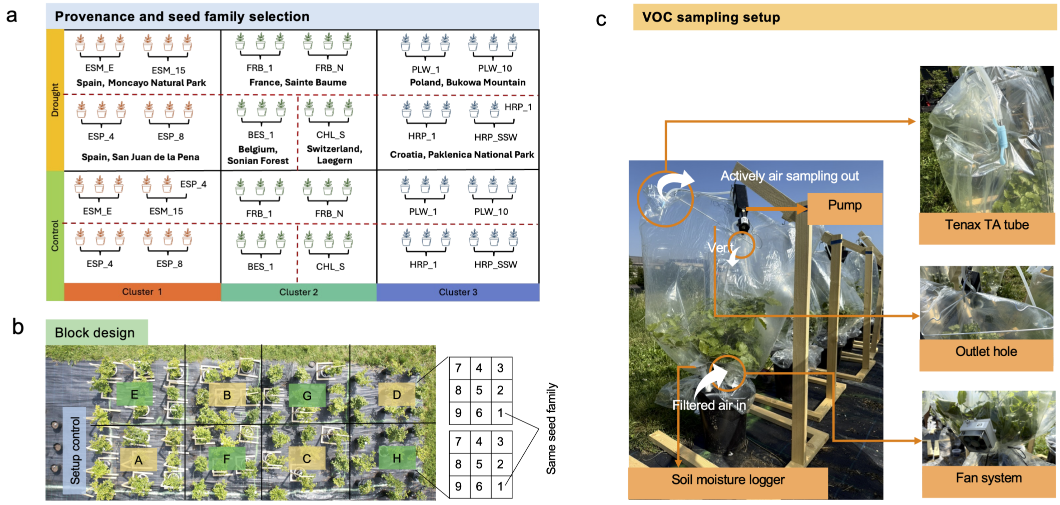


**Figure S1. Overview of the genetic variation source, experimental design, and VOC sampling setup.** (a) Genetic clusters, provenances, and seed families used in this study; the country and the locality of the provenances were indicted by bold text (b) experimental layout; and (c) VOC sampling setup.

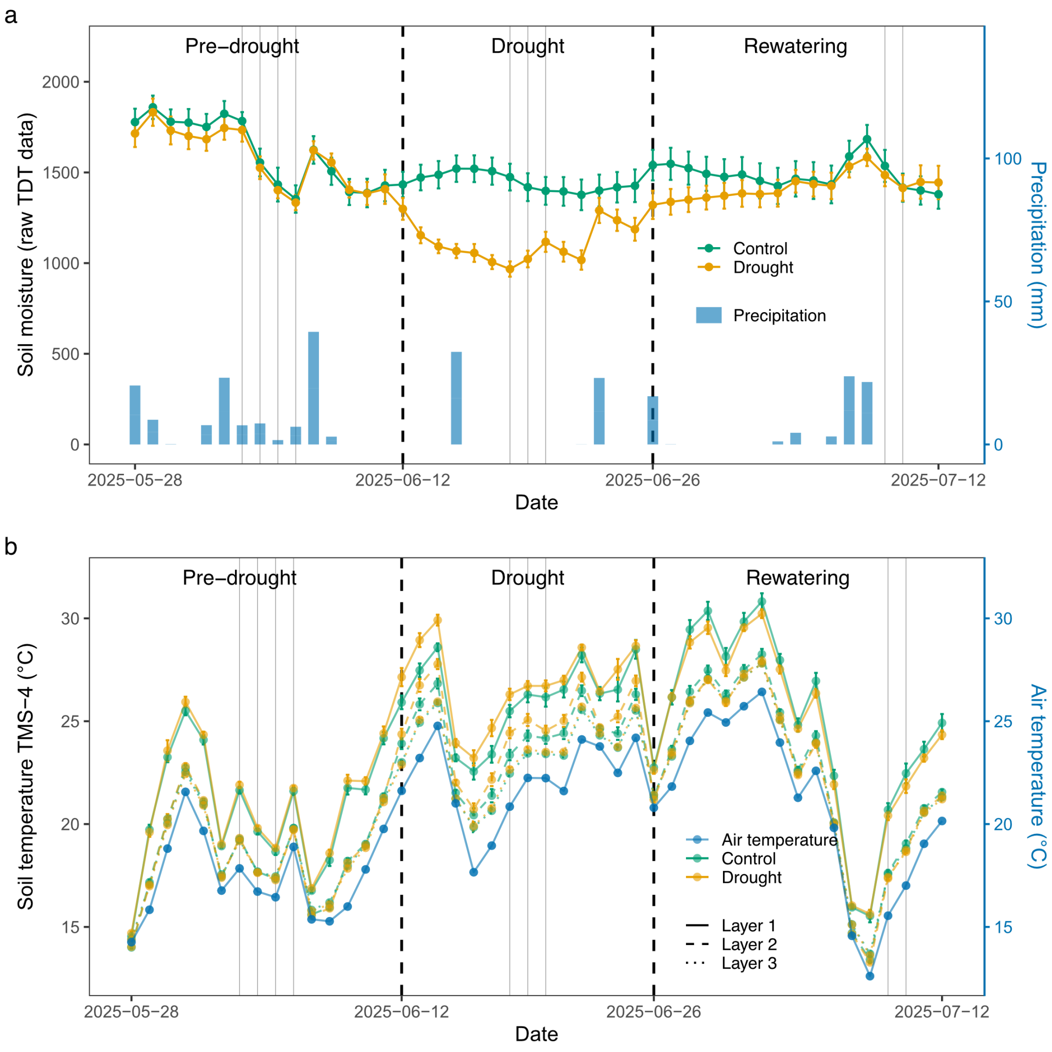


**Figure S2. Environmental moisture and temperature during the experiment** (a) Soil moisture and precipitation. Soil moisture are shown as mean ± SE. Precipitation represents daily total precipitation. (b) soil and air temperature during the three experimental periods. and soil temperature (measured at three depths: layer 1: +15 cm, Layer 2: 0, and layer 3: −8 cm relative to the soil surface). Air temperature represents daily mean temperature. VOC sampling dates are indicated by gray vertical lines. Control group is indicated by green and drought group is indicated by yellow.

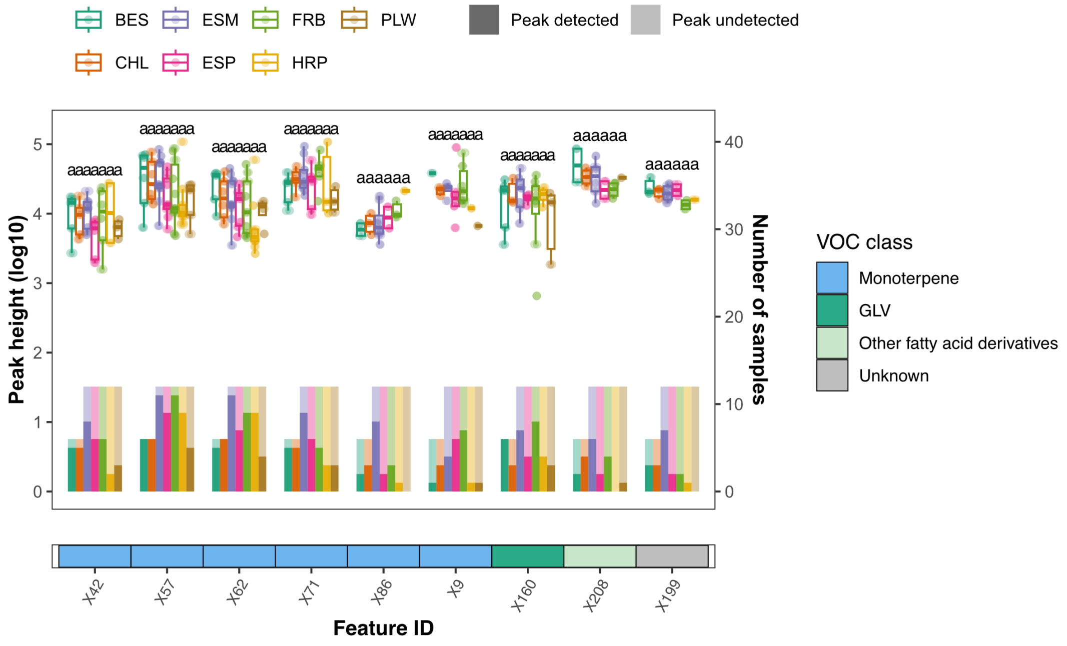


**Figure S3. Peak heights of significant VOC features across the seven provenances.** Each column represents a VOC feature, and its classification is indicated by the colored bar below the x-axis. Box plots show the distribution of peak heights for samples in which the feature was detected, with individual sample values overlaid as scatter points. The lower, middle, and upper hinges correspond to the first, median, and third quartiles (25th, 50th, and 75th percentiles), and whiskers extend to 1.5 × the interquartile range. Bars below each box plot indicate the number of tree samples in which the feature was detected (dark color) or not detected (light color). Provenances are distinguished by color. Different lowercase letters above the box plots and bars indicate significant differences among provenances based on Tukey's HSD test ($\alpha = 0.05$). Bars without letters indicate insufficient sample size to fit a generalized linear model.

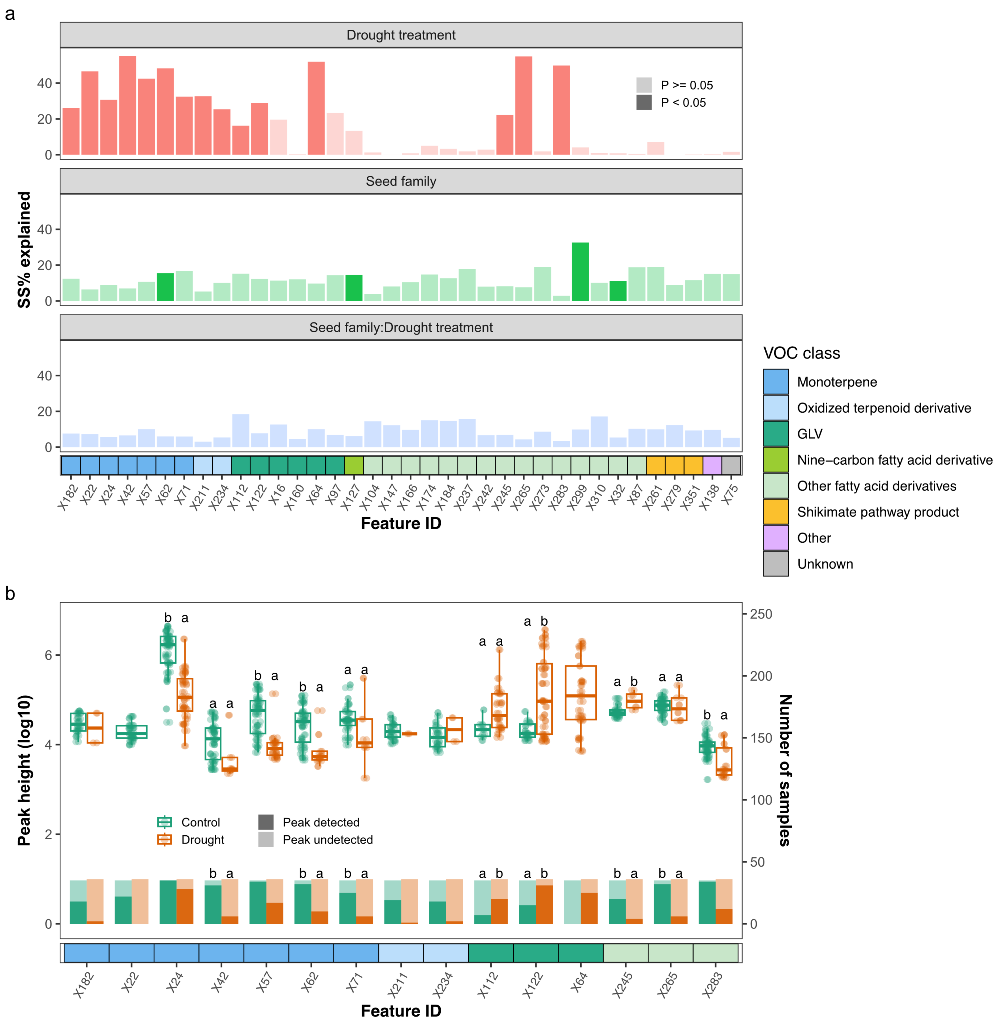


**Figure S4. Drought treatment effects on VOC variation during the drought period.** (a) Variation in VOC features explained by drought treatment, seed family, and their interaction. Drought treatment, seed family, and their interaction were fitted sequentially in ANOVAs to calculate the percentage of the sum of squares (SS%), representing the proportion of variance explained by each factor. Significance is indicated by bar color (dark: $P < 0.05$; light: $P \geq 0.05$). (b) peak heights and detection frequencies of VOC features showing significant drought treatment effects. Each column represents a VOC feature, and its classification is indicated by the colored bar below the x-axis. Box plots show the distribution of peak heights for samples in which each feature was detected. The lower, middle, and upper hinges represent the first, median, and third quartiles (25th, 50th, and 75th percentiles), and whiskers extend to $1.5 \times$ the interquartile range. Bars below each box plot indicate the number of tree samples in which the feature was detected (dark color) or not detected (light color). Control and drought groups are distinguished by color. Different lowercase letters above the box plots and bars indicate significant differences between treatments based on Tukey's HSD test ($\alpha = 0.05$). Box plots and bars without letters indicate insufficient sample size to fit linear or generalized linear models.

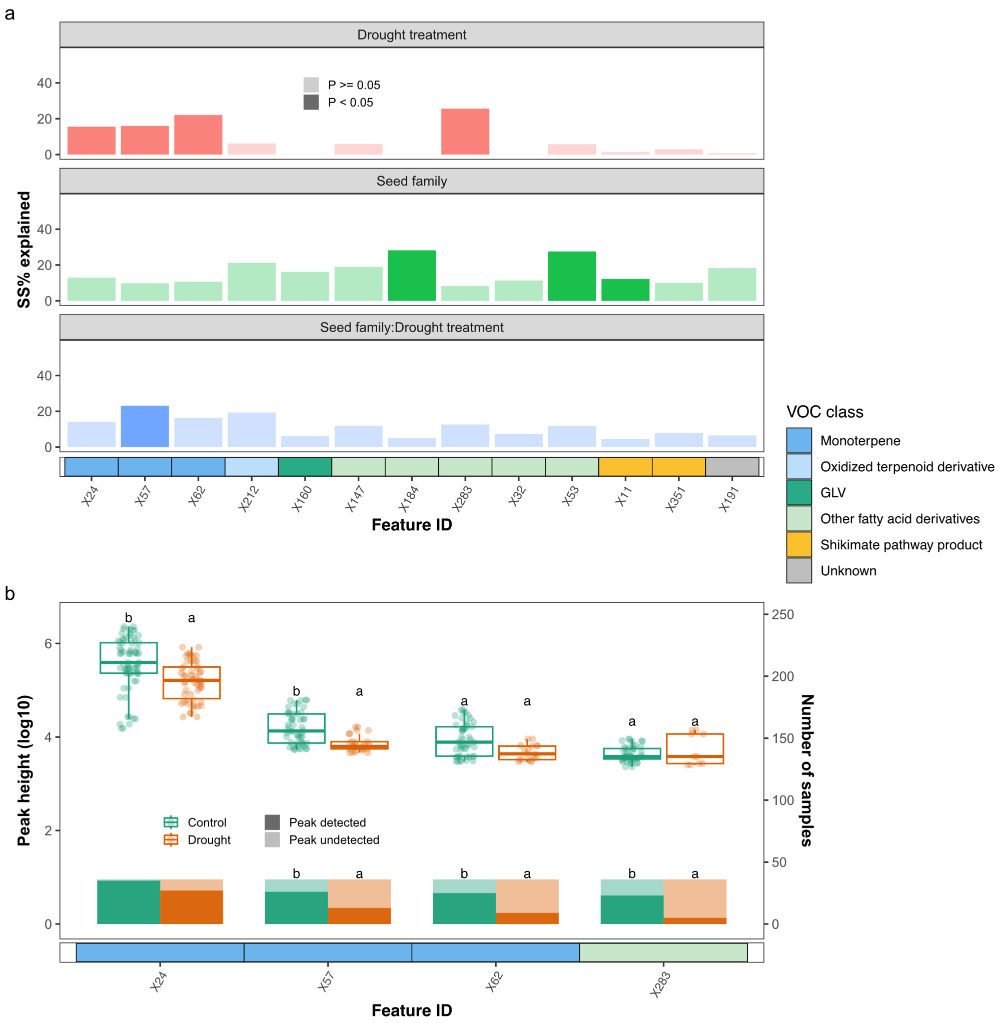


**Figure S5. Drought treatment effects on VOC variation during the rewatering period.** (a) Variation in VOC features explained by drought treatment, seed family, and their interaction. Drought treatment, seed family, and their interaction were fitted sequentially in ANOVAs to calculate the percentage of the sum of squares (SS%), representing the proportion of variance explained by each factor. Significance is indicated by bar color (dark: $P < 0.05$; light: $P \geq 0.05$). (b) peak heights and detection frequencies of VOC features showing significant drought treatment effects. Each column represents a VOC feature, and its classification is indicated by the colored bar below the x-axis. Box plots show the distribution of peak heights for samples in which each feature was detected. The lower, middle, and upper hinges represent the first, median, and third quartiles (25th, 50th, and 75th percentiles), and whiskers extend to $1.5 \times$ the interquartile range. Bars below each box plot indicate the number of tree samples in which the feature was detected (dark color) or not detected (light color). Control and drought groups are distinguished by color. Different lowercase letters above the box plots and bars indicate significant differences between treatments based on Tukey's HSD test ($\alpha = 0.05$). Box plots and bars without letters indicate insufficient sample size to fit linear or generalized linear models.
